# Mouse behavioral genomics identifies Creld1 as a gatekeeper of somatosensation

**DOI:** 10.64898/2026.03.30.715210

**Authors:** Yan Jiang, Wanru Huang, Hailong Yao, Yan Chen, Cheng Bi, Tianyuan Ye, Shangkun Wang, Hongda Yin, Bailong Xiao

**Author notes:** To whom correspondence should be addressed: Bailong Xiao. These authors contribute equally to this work (Yan Jiang, Wanru Huang).

## Abstract

Somatosensation enables the perception of touch, temperature, pain, and itch. These sensory modalities are mediated by primary sensory neurons in the dorsal root ganglion (DRG), which convert physical and chemical stimuli into electrical signals using specialized molecular sensors, including the touch sensor PIEZO2 (Ref^1,2^) and temperature sensors such as TRPV1 (Ref^3,4^), coupled with downstream voltage-gated sodium channels (Na_v_) like Na_v_1.7 (Ref^5,6^). However, the full repertoire of molecular components involved in somatosensory processing remains incompletely identified. Here, we developed an efficient postnatal CRISPR-Cas9 knockout platform to screen DRG-expressed genes via AAV9-sgRNA delivery. Combining this approach with behavioral assays of somatosensory responses, we validated the roles of PIEZO2 and TRPV1 in sensing gentle touch and noxious heat, respectively, and the broad involvement of Na_v_1.7 in distinct somatosensory modalities. Remarkably, a targeted screen of 20 DRG-expressed genes identified the Cysteine-rich with EGF-like domains 1 (Creld1) as a master regulator of somatosensation. Either sgRNA-mediated postnatal knockout or tamoxifen-induced Cre-mediated deletion of Creld1 in DRG neurons resulted in profound behavioral deficits in touch, temperature, pain, and itch perception, while its overexpression enhanced touch and thermal responses. Mechanistically, Creld1 functions as a novel auxiliary regulator of Na_v_ for controlling the excitability of DRG neurons. The biochemical interaction and functional modulation of Na_v_ by mouse Creld1 are mediated via its C-terminal transmembrane region. Notably, this domain is conserved in mouse Creld1 and some isoforms of human CRELD1, resulting in an isoform-dependent regulation of Na_v_1.7. Together, this work establishes a robust postnatal screening platform for somatosensory genomics and identifies Creld1 as a master regulator of somatosensory function, and provides novel therapeutic strategy for pain and itch treatment.

## Main

DRG neurons are specialized primary sensory neurons that detect and transduce chemical, thermal, and mechanical stimuli into action potentials dictated by voltage-gated sodium channel (VGSC) isoforms, such as Na_v_1.6, Na_v_1.7, and Na_v_1.8 (Ref^7^). Reflecting their varied sensory roles, DRG neurons exhibit significant heterogeneity in morphology, electrophysiological properties, and molecular expression profiles^8^. To systematically elucidate DRG-mediated somatosensation—encompassing touch, temperature, pain, itch—extensive research has focused on identifying and characterizing the molecular receptors and regulators involved. Over the past three decades, diverse genotype-phenotype screening strategies have been employed. For example, cloning and heterologous expression of DRG-derived cDNA libraries led to the discovery of TRPV1 as the receptor for capsaicin and noxious heat and TRPM8 as the receptor for menthol and cool temperature^3,4,9,10^. Alternatively, targeted siRNA knockdown combined with patch-clamp recording of mechanically activated currents identified PIEZO2 in the mechanosensitive PIEZO channel family as the touch receptor in DRG neurons^1,11^. Additionally, genomic deletion of candidate genes, such as the G protein-coupled receptor family Mrgprs, revealed MrgprA3 as the chloroquine-sensitive itch receptor^12^. These findings have significantly advanced our understanding of the molecular and cellular mechanisms underlying temperature, touch, pain, and itch sensation.

Despite this progress, the full repertoire of molecular receptors and regulatory factors governing somatosensory function remain incompletely characterized. Traditional knockout approaches face limitations due to high costs, low throughput, developmental compensation, or lethality, complicating the study of DRG-expressed genes in postnatal somatosensory functions. To overcome these challenges, we developed an innovative and efficient postnatal gene knockout strategy targeting DRG neurons. This approach enables systematic in vivo behavioral genomic screening of candidate genes and has identified Creld1 as a master regulator of somatosensory behaviors.

## Results

### Postnatal gene knockout in DRG neurons using the CRISPR-Cas9 system

To enable targeted genomic screening of somatosensory behaviors, we developed an efficient postnatal DRG gene knockout strategy using CRISPR-Cas9. We intracerebroventricularly injected AAV9 virus carrying the single-guide RNAs (sgRNAs) targeting the gene of interest and the green fluorescent protein (GFP) marker into Cas9-expressing mice at postnatal day 0–2 (P0–P2) (Fig. 1a). The virus infection efficiency was assessed by immunostaining GFP and the markers for labeling distinct subpopulations of DRG neurons or other tissues, including the neuronal maker TUBB3 (βIII-tubulin), the Low Threshold Mechanoreceptor (LTMR) and proprioceptor marker NFH (Neurofilament Heavy Chain), the non-peptidergic nociceptor marker IB4, and the peptidergic nociceptor marker CGRP (Calcitonin Gene-Related Peptide). GFP fluorescence staining revealed sparse infection in the brain, spinal cord, and footpad skin (Extended Data Fig. 1a), but more abundant infection in vagus ganglion neurons and DRG neurons (Extended Data Fig. 1a, b). The immunostaining result in DRG neurons revealed efficient infection across DRG neuron subpopulations, with GFP^+^ neurons detected in 78.5 ± 2.0% of TUBB3^+^, 72.0 ± 3.2% of IB4^+^, 77.5 ± 4.0% of CGRP^+^, and 71.8 ± 4.7% of NFH^+^ neurons (Extended Data Fig. 1b, c). The proportions of IB4^+^ (29.2 ± 1.8%), CGRP^+^ (21.8 ± 1.8%), and NFH^+^ (43.8 ± 5.1%) neurons among GFP^+^ cells matched their natural distributions (Extended Data Fig. 1b, d). Thus, these data demonstrate effective and unbiased infection of DRG neurons by the intracerebroventricularly injected AAV9 virus.

**Fig. 1.**
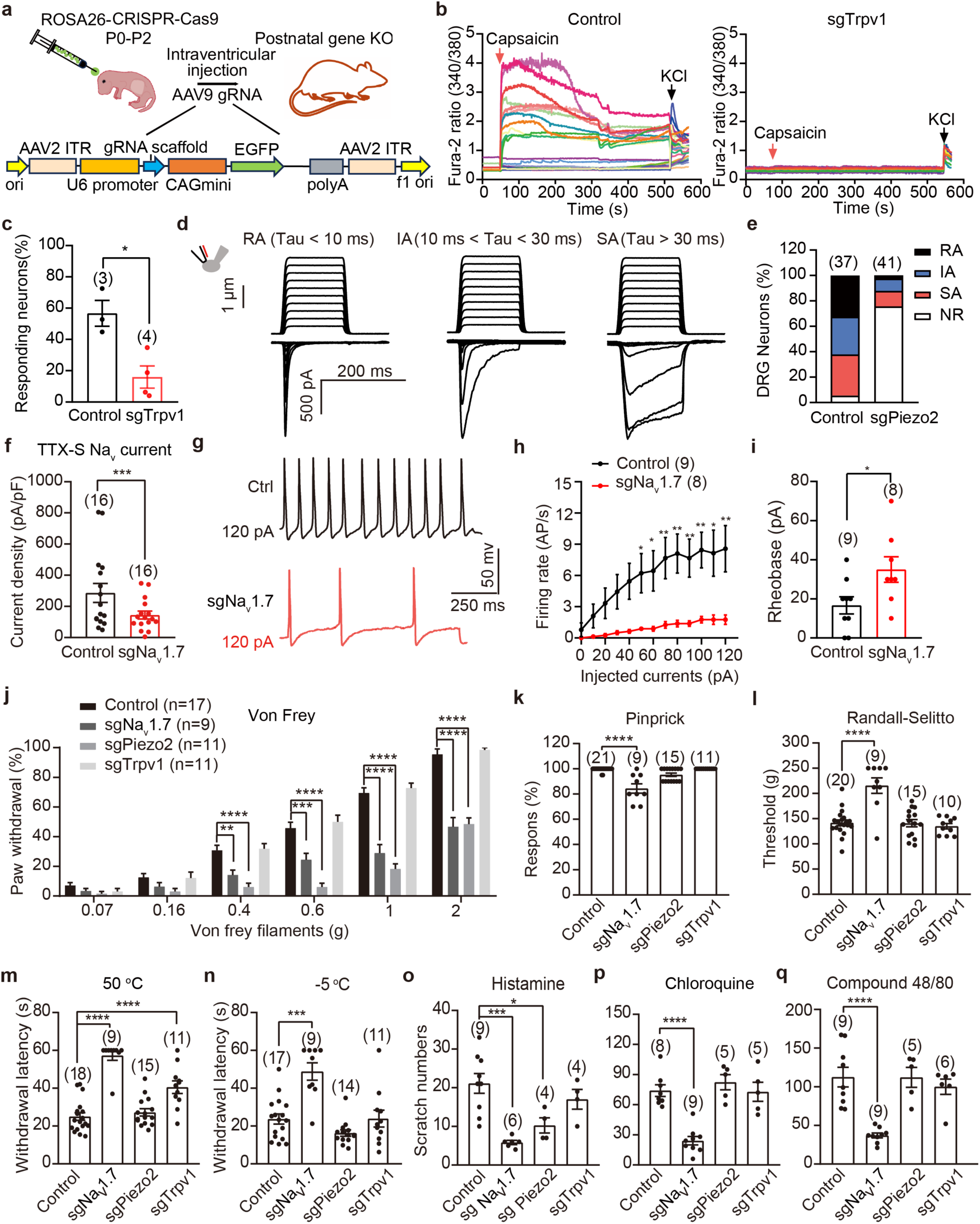
Postnatal knockout of DRG-expressed genes. **a**, Schematic diagram of the CRISPR/Cas9-mediated knockout of target genes expressed in DRG neurons via intraventricular injection of the AAV9 virus carrying the construct containing the sgRNA against the target gene and EGFP for indication of infection efficiency. 6-8 weeks post injection, the mice were subjected to experimental tests. **b**, Representative single-cell Ca^2+^ imaging of DRG neurons derived from control sgRNA- or sgTrpv1-targeted mice in response to 10 μM capsaicin and 50 mM KCl. Individual neurons are color-coded. **c**, Scatterplot of the percentage of cultured GFP^+^ DRG neurons showing capsaicin-induced Ca^2+^ response. **d**, Representative traces of mechanically evoked whole-cell currents of rapidly inactivating (RA), intermediate inactivating (IA), and slowly inactivating (SA) kinetics recorded from cultured control DRG neurons. **e**, Proportion of control sgRNA- and sgPiezo2-targted GFP^+^ DRG neurons showing the indicated responding properties. **f**, The peak current density of tetradotoxin (TTX)-sensitive Na_v_ current of DRG neurons derived from the control sgRNA-and sgNa_v_1.7-targeted mice. **g**, Representative trace of evoked action potential (APs) of the control sgRNA- and sgNa_v_1.7-targeted DRG neurons. **h**, Firing frequency of the control sgRNA- and sgNa_v_1.7-targeted DRG neurons in response to the injected currents. **i**, Rheobase of sgNa_v_1.7-targeted DRG neurons compared with the control sgRNA-targeted neurons. **j**, The percentage of paw withdrawals in response to the indicated series of Von Frey filament stimulation. **k**, Scatterplot of the percentage of paw withdrawal in response to the pinprick test. **l**, Scatterplot of the threshold in response to the Randall-Selitto test. **m**, Scatterplot of the paw withdrawal latency in response to the hot plate test. **n**, Scatterplot of the paw withdrawal latency in response to the cold plate test. **o-q**, Scatterplot of the scratch numbers within 30 minutes in mice injected with histamine (**o**), chloroquine (**p**) or compound 48/80 (**q**). Data are presented as means ± SEM; sample sizes are indicated. Statistical significance is determined by Unpaired Student’s *t* test for **c**, **f**, **i**. Two-way ANOVA with Bonferroni’s multiple comparisons test for **j**. One-way ANOVA with Bonferroni’s multiple comparisons test for **k**-**q**, \**P* < 0.05, \*\**P* < 0.01, \*\*\**P* < 0.001, \*\*\*\**P* < 0.0001.

We validated the knockout efficiency using Trpv1, Piezo2, and Na_v_1.7 as representative targets. Eight weeks post-injection of their respective AAV-sgRNAs (sgTrpv1, sgPiezo2 and sgNa_v_1.7) (Extended Data Table 1), quantitative PCR showed mRNA reductions of 55.8% for Na_v_1.7, 51.7% for Piezo2, and 70.1% for Trpv1 compared to controls (Extended Data Fig. 1f-h). Furthermore, the mRNA levels of non-target genes were not reduced (Extended Data Fig. 1f-h). Targeting Na_v_1.7 did not affect the expression of other Na_v_ members including Na_v_1.6, Na_v_1.8 and Na_v_1.9 (Extended Data Fig. 1i). These data indicate no substantial off-target effects. Immunofluorescent staining with an anti-Na_v_1.7 antibody revealed apparent expression and localization of the Na_v_1.7 proteins in the plasma membrane of DRG neurons derived from control mice, which was markedly decreased in DRG neurons derived from sgNa_v_1.7-targed mice (Extended Data Fig. 1j). Furthermore, due to the lack of highly specific antibody suitable for immunostaining of Piezo2 and Trpv1, we carried out hybridization chain reaction staining and found drastically reduced mRNA expression of Piezo2 in sgPiezo2-targeted DRG neurons and Trpv1 in sgTrpv1-targeted DRG neurons, respectively (Extended Data Fig. 1k, l).

Consistent with the expression analyses, functional assessments demonstrated specific deficits corresponding to each knockout. Trpv1 mediates capsaicin-induced Ca^2+^ response in DRG neurons. Using single-cell Ca^2+^ imaging, we found that Trpv1 knockout reduced the percentage of capsaicin-responsive neurons from 56.7 ± 8.2% in controls to 16.0 ± 7.1% (Fig. 1b, c). Using whole-cell patch clamp equipped with piezo-driven glass pipette for cell indentation, we recorded mechanically activated currents in control DRG neurons with distinct inactivation kinetics (Fig. 1d), which were classified as rapidly adapting (RA), intermediately adapting (IA), or slowly adapting (SA). In line with that Piezo2 predominantly mediates the RA current^1^, the proportion of RA neurons was markedly decreased in the sgPiezo2-targeted mice (Fig. 1e). Na_v_1.7 in DRG neurons mediates tetradotoxin (TTX)-sensitive Na_v_ current^13^. Na_v_1.7 knockout resulted in reduced TTX-sensitive currents, decreased action potential firing, and increased rheobase (Fig. 1f-i, Extended Data Fig. 1e), confirming the crucial role of Na_v_1.7 in affecting the excitability of DRG neurons.

To examine whether postnatal knockout of Trpv1, Piezo2 and Na_v_1.7 could lead to behavioral deficits of their somatosensory functions, we conducted a battery of behavioral tests, including Von Frey for gentle touch (Fig. 1j), pinprick (Fig. 1k) and Randall-Selitto (Fig. 1l) for mechanical pain, 50 °C hot plate for noxious heat (Fig. 1m), -5 °C cold plate for noxious cold (Fig. 1n), and pruritogen (histamine-dependent and histamine-independent chloroquine and Compound 48/80)-induced itch (Fig. 1o-q). The sgPiezo2-targeted mice exhibited nearly abolished responses to Von Frey filaments below 0.6 g of forces (Fig. 1j), an effect even stronger than those observed in the Piezo2-cKO mice driven by the Advillin-Cre-mediated deletion of Piezo2 in a large set of DRG neurons and in Merkel cells^2^. The sgPiezo2-targeted mice appeared normal in response to noxious mechanical or temperature stimuli (Fig. 1k-n), in line with its specific role in sensing gentle touch, but not acute mechanical pain and temperature^2,14^. Notably, the sgPiezo2-targeted mice showed impaired response to histamine-induced itch, but normal response to chloroquine- or compound 48/80-induced itch (Fig. 1o-q). Consistent with the role of Trpv1 in sensing noxious heat^4^, sgTrpv1-targeted mice showed defective responses only in noxious heat sensitivity (Fig. 1m). Global deletion of Na_v_1.7 in C57BL/6 mice was originally reported with lethality shortly after birth, preventing behavioral analyses in adult mice^5^. Thus, the role of Na_v_1.7 in distinct sensory modalities has been systematically investigated using different Cre mouse lines, including Na_v_1.8-Cre, Advillin-Cre, Wnt-Cre and tamoxifen-inducible ERT-Cre^5,15–19^. These extensive characterizations have demonstrated the critical role of Na_v_1.7 in transducing mechanical, thermal and chemical pain as well as chloroquine- or histamine-induced itch^18–20^. Using genetic and animal husbandry strategies to overcome the neonatal-lethal phenotype, global Na_v_1.7 knockout mice in CD1 outbred strain were obtained and their behavioral characterizations verified the critical role of Na_v_1.7 in noxious mechanical and heat sensation^21^. Remarkably, sgNa_v_1.7-targeted mice in C57BL/6 background were viable and displayed broad and severe impairments of noxious mechanical, heat and cold responses as well as histamine-, chloroquine-, compound 48/80-induced itch (Fig. 1k-o), which recapitulates phenotypes previously observed in the conditional or global Na_v_1.7 knockout mice^5,15–17,20,21^. Notably, sgNa_v_1.7-targeted mice showed significantly defective responses to Von Frey filaments at 0.4-2 g of forces, which is similar to the sgPiezo2-targed mice (Fig. 1j). These data suggest that Na_v_1.7 might also be involved in gentle touch sensation, which is consistent with the finding that Na_v_1.7 is expressed in C-low threshold mechanoreceptors (C-LTMRs) and might mediate affective touch in both mice and human^22^. Collectively, these results validate our postnatal knockout strategy for functional analysis of DRG genes in adult somatosensory functions, which might largely overcome embryonic or perinatal lethality and developmental compensation associated with conventional embryonic knockout approach.

### Targeted genomics screen identifies Creld1 as a pan-sensory regulator

To demonstrate the utility of our postnatal DRG gene knockout platform for identifying novel molecular regulators of somatosensation, we performed a targeted behavioral genomic screen of 20 candidate genes, which were roughly classified as (1) DRG subtype classification markers^23^: Pvalb (Parvalbumin), Bmpr1b (Bone morphogenetic protein receptor type-1B), Ret (Proto-oncogene tyrosine-protein kinase receptor), Sstr2 (Somatostatin receptor type 2), Th (Tyrosine 3-monooxygenase), Ntrk3 (NT-3 growth factor receptor), Adra2a (Alpha-2A adrenergic receptor), Ntrk2 (BDNF receptor), Tac1 (Protachykinin-1), and Calca (Calcitonin); (2) DRG-expressed membrane proteins including ion channels, receptors, transporters: Chrna9 (Neuronal acetylcholine receptor subunit alpha-9), Stoml3 (Stomatin-like protein 3), Mrgprg (Mas-related G-protein coupled receptor member G), Scarb2 (Lysosome membrane protein 2), Vdac1 (Non-selective voltage-gated ion channel 1), Vdac3 (Non-selective voltage-gated ion channel 3), Slc44a2 (Choline transporter-like protein 2), Nnt [NAD(P) transhydrogenase), Pannexin1 and Creld1 (Cysteine-rich with EGF-like domains 1) (Fig. 2a).

**Fig. 2.**
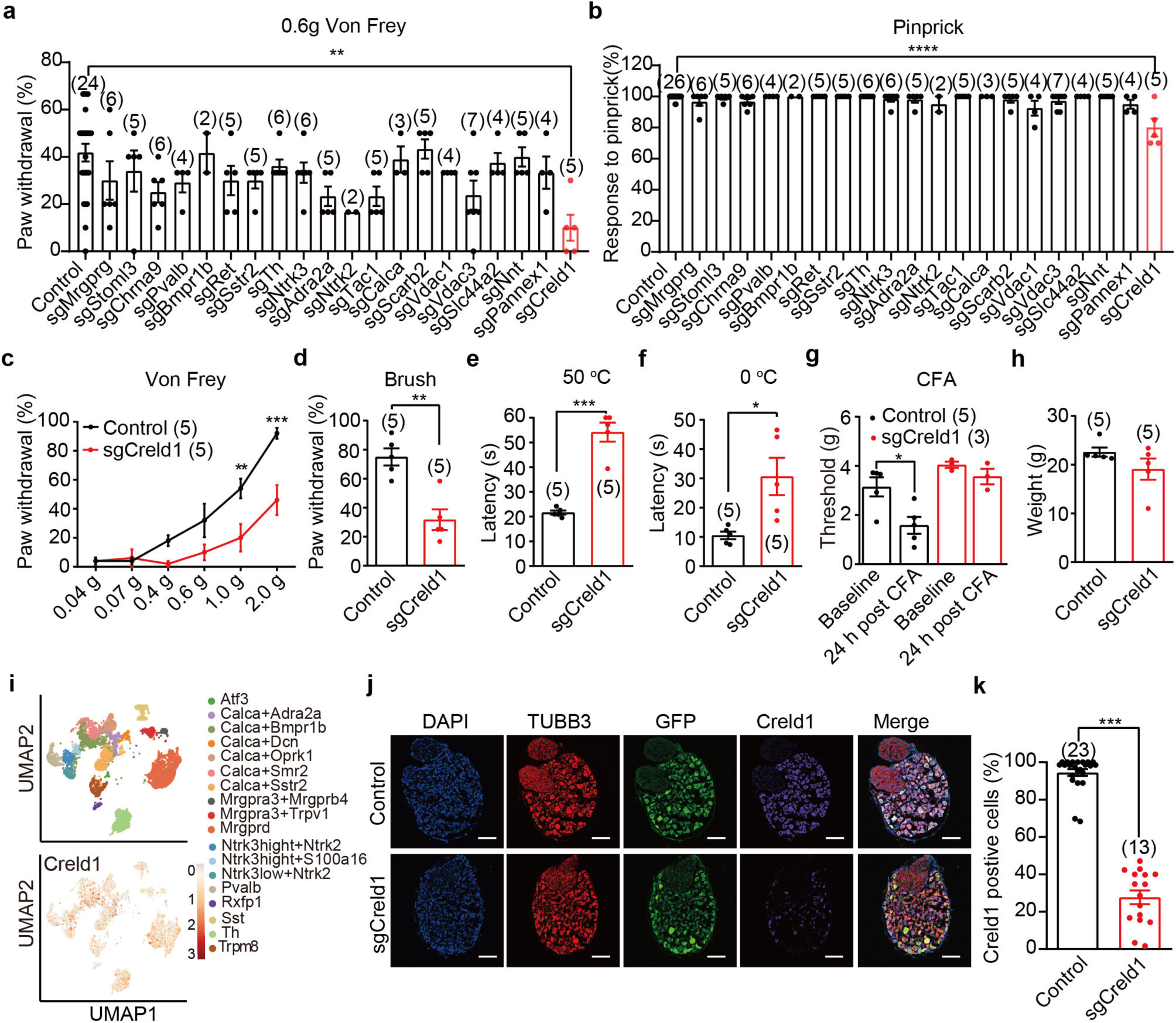
Behavioral genetics screening identifies Creld1 as a critical regulator of somatosensory behaviors. **a** and **b**, Scatterplots of the percentage of paw withdrawal in response to 0.6 g Von Frey filament (**a**) or pinprick (**b**) stimulation of mice injected with the indicated sgRNAs against the targeted genes. **c**, The percentage of paw withdrawal in response to the indicated series of Von Frey filament stimulation. **d**, Scatterplot of the percentage of paw withdrawal in response to brush. **e** and **f**, Scatterplot of the paw withdrawal latency in response to the hot plate (**e**) and cold plate (**f**). **g**, Scatterplot of the paw withdrawal threshold in the ramping Von Frey test of baseline (before CFA injection) and 24 h post CFA injection in control and sgCreld1 female mice. **h**, Scatterplot of the body weight. **i**, Top: UMAP projection of harmonized neuronal atlas. Cells are colored by the annotated cell types that are defined by the indicated marker genes. Bottom: Creld1 expression across DRG neuron subtypes. Color intensity indicates mean log-normalized expression values. **j**, Representative images of immunostaining of DRG sections from control and sgCreld1 mice. Scale bar: 50 μm. **k**, Scatterplot of the percentage of Creld1 positive cells in control and sgCreld1 DRG neurons. Data are presented as means ± SEM; sample sizes are indicated. One-way ANOVA with Bonferroni’s multiple comparisons test for **a**, **b**. Two-way ANOVA with Bonferroni’s multiple comparisons test for **c**, and unpaired Student’s *t* test for **d**-**h**, **k**, \**P* < 0.05, \*\**P* < 0.01, \*\*\**P* < 0.001.

AAV9 viruses encoding individual sgRNAs (Extended Data Table 1) were intracerebroventricularly injected into Cas9-expressing mice at P0-P2. Notably, only 2 mice per group survived to adulthood in the sgBmpr1b and sgNtrk2 targeted cohorts. At eight weeks post-injection, we assessed body weight and conducted comprehensive behavioral testing, including Von Frey, pinprick, Randall-Selitto, hot plate, and cold plate assays. All sgRNA-targeted groups showed comparable body weights to control sgRNA-targeted mice (Extended Data Fig. 2a), indicating no gross developmental abnormalities. Unexpectedly, postnatal knockout of the candidate genes previously implicated in somatosensory function such as Chrna9^24^ and Stoml3^25^ or DRG classification did not cause detectable behavioral deficits (Fig. 2a, b and Extended Data Fig. 2b-d). For example, deletion of the nociceptor marker gene Tac1 (encoding the substance P, SP) or Calca (encoding the calcitonin gene related peptide, CGRP) did not cause behavioral defects (Fig. 2a, b and Extended Data Fig. 2b-d). The results are consistent with no behavioral defects observed in mice lacing both Tac1 and Caca^26^. Stoml3 was reported to be essential for touch sensation^25^. In contrast, we did not observe touch deficit in sgStoml3-targeted mice (Fig. 2a, b and Extended Data Fig. 2b-d). Quantitative PCR analysis revealed a significant reduction in the mRNA expression level of the target gene in the DRG of mice injected with the target-specific gRNA, relative to the control group, with expression levels reduced by 74.9% for Mrgprg, 50.9% for Stomls, 68.8% for Chrna9, 69.7% for Pvalb, 75.7% for Bmpr1b, 59.6% for Ret, 71.1% for Sstr2, 77.4% for Th, 87.9% for Ntrk3, 59.7% for Adra2a, 41.9% for Ntrk2, 50.2% for Tac1, 70.9% for Calca, 44.0% for Scarb2, 69.8% for Vdac1, 77.6% for Vdac3, 52.1% for Slc44a2, 77.6% for Nnt and 77.0% for Pannex1 (Extended Data Fig. 2e). Given that the mRNA reduction of the candidate genes was largely within the range of ∼50-70% reduction detected in sgNa_v_1.7-, sgPiezo2-, or sgTrpv1-targeted mice (Extended Data Fig. 1f-h), we considered that the lack of behavioral deficits might not be due to knockout insufficiency. In light of the prominent role of Piezo2 in sensing touch (Fig. 1j), we reasoned that the previously observed touch deficits of the Stoml3 KO mice might alternatively be due to developmental effect^25^.

In striking contrast to the other groups, the sgCreld1-targeted mice exhibited profound deficits across all tested somatosensory modalities. These animals showed significantly impaired responses to light touch of 0.6 g of Von Frey (Fig. 2a) and noxious mechanical stimulation of pinprick (Fig. 2b). Follow-up behavioral characterizations of the sgCreld1-targted mice confirmed and extended the initial screening findings. The sgCreld1 group showed deficits in response to a serial of Von Frey filaments ranging from 0.04 to 2.0 g (Fig. 2c), brush (Fig. 2d), hot plate (Fig. 2e) and cold plate (Fig. 2f). We further tested the involvement of Creld1 in Complete Freund’s adjuvant (CFA)-induced inflammatory pain model (Fig. 2g). CFA-treatment induced a significant reduction of the mechanical pain threshold in control group (Fig. 2g). By contrast, such CFA-induced mechanical allodynia was not observed in the sgCreld1-targted group (Fig. 2g). The sgCreld1-targted mice had normal weight at 5 - 7 weeks (Fig. 2h) and rotarod performance (Extended Data Fig. 2f). These findings suggest that the observed defective somatosensory responses might not be due to gross defects of motility or fitness. Analyses of single-cell RNA sequencing data^27^ revealed widespread distribution of Creld1 across molecularly distinct DRG neuron subtypes (Fig. 2i). Immunostaining using an anti-Creld1 antibody confirmed abundant Creld1 protein expression in control DRG neurons, while showing marked reduction in sgCreld1-targeted neurons (Fig. 2j, k). Efficient knockout of Creld1 in the sgCreld1-targeted mice was also evidenced by 57.8% reduction in the mRNA level and 77.1% decrease in the protein expression (Extended Data Fig. 2g-i). Collectively, these findings identified Creld1 as a robust regulator of somatosensory functions.

### Conditional knockout of Creld1 in DRG neurons recapitulates the somatosensory defects

Creld1 was originally cloned and characterized as a new member of the epidermal growth factor (EGF) superfamily and showed abundant expression in human heart and brain tissues^28^. Subsequent studies have found that Creld1 functions in cardiac development and immune regulation in both mouse and human, with mutations associated with atrioventricular septal defects (AVSD) and neurodevelopmental effects including early-onset epilepsy^29–32^. Our findings reveal its significant role in sensory neurons. RT-PCR analysis showed prominent Creld1 expression in DRG neurons, exceeding levels detected in heart, brain and skin tissues (Extended Data Fig. 3a). To determine whether the behavioral deficits resulted specifically from loss of DRG-expressed Creld1 rather than indirect effects from other tissues, we generated Creld1 floxed mice with loxP sites flanking exons 3-4 (Creld1^fl/fl^) (Extended Data Fig. 3b, c). When crossed to several commonly used DRG-specific Cre lines including Piezo2-Cre^2^, Pirt-Cre^33^ and Advillin-Cre^34^, the resulting Piezo2-Cre/Creld1^fl/fl^, Pirt-Cre/Creld1^fl/fl^ and Advillin-Cre/Creld1^fl/fl^ mice were embryonic lethal. While the exact cause of the lethality remains to be determined, these observations demonstrate the critical role of Creld1 in the primary sensory neurons. To assay the role of Creld1 in adult somatosensory functions, we crossed the Creld1^fl/fl^ mice with the Advillin-CreERT2 mice^35^ and achieved tamoxifen-inducible, DRG-specific deletion of Creld1 (cKO). Following tamoxifen administration at two weeks, quantitative analysis revealed a 90.8% reduction in Creld1 transcript and 81.7% decrease in protein levels in DRG neurons of cKO mice compared to controls (Extended Data Fig. 3d-f). Immunostaining confirmed specific reduction of Creld1 in DRG neurons (Extended Data Fig. 3g, h).

When subjected to the battery of behavioral assays, the cKO mice displayed significant somatosensory impairments across multiple behavioral modalities (Fig. 3), closely recapitulating the deficits observed in the sgCreld1-targeted mice (Fig. 2). Specifically, the cKO mice showed reduced sensitivity to touch when tested with Von Frey filaments (Fig. 3a) and brush stimulation (Fig. 3b). Their responses to noxious mechanical stimuli were also diminished, as evidenced by decreased sensitivity in both pinprick (Fig. 3c) and Randall-Selitto tests (Fig. 3d). Thermal nociception was impaired, with attenuated responses to both heat and cold stimuli (Fig. 3e, f). Moreover, the cKO mice also lost capsaicin-induced mechanical allodynia (Fig. 3g). The remarkable similarity between the behavioral phenotypes of cKO and sgCreld1-targeted mice strongly supports a DRG-specific role for Creld1 in mediating diverse somatosensory modalities.

**Fig. 3.**
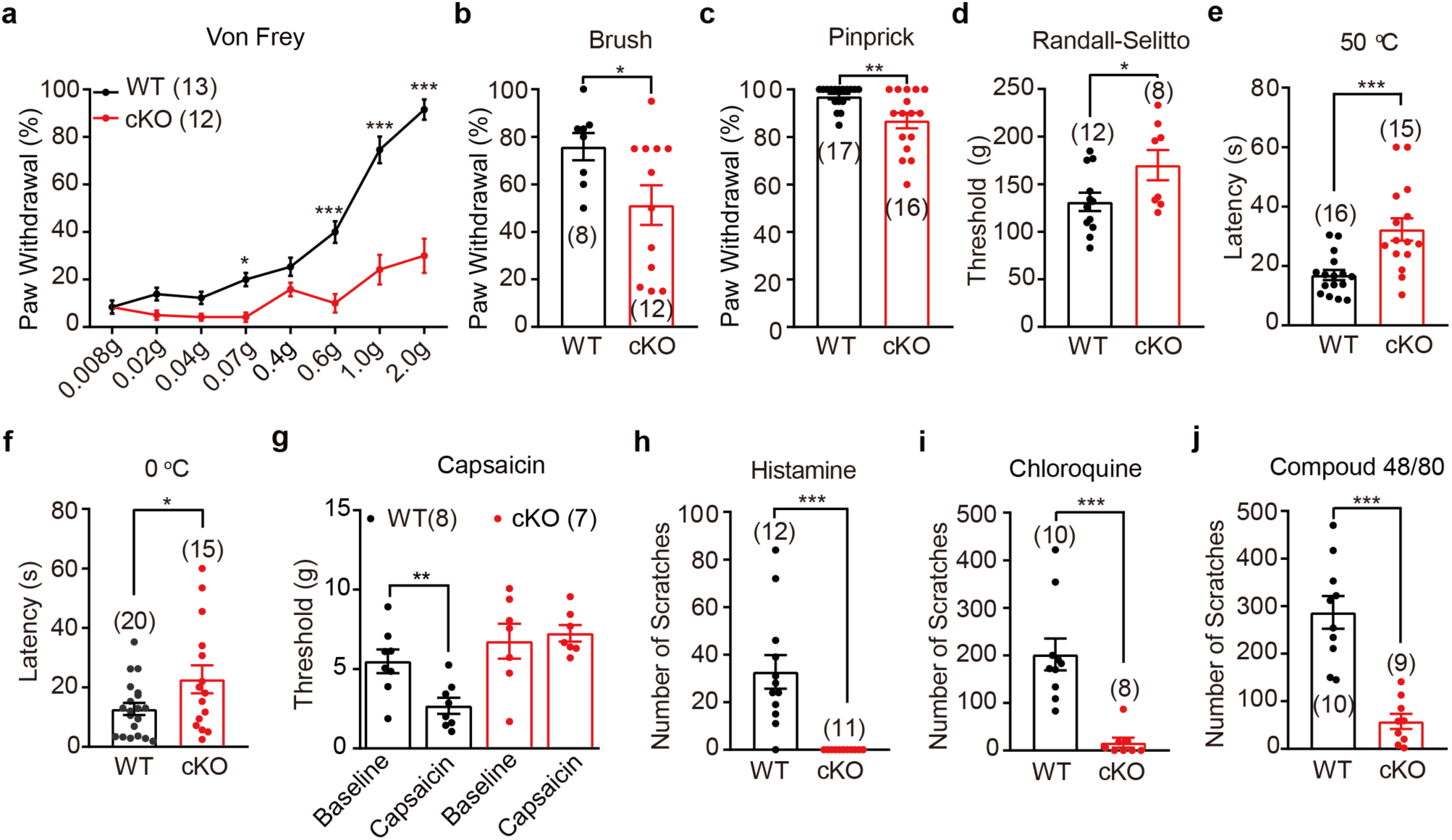
DRG-specific knockout of Creld1 impairs multiple somatosensory modalities. **a**, **b**, The paw withdrawal response of the control and cKO mice (Advillin-CreERT2/Creld1^fl/fl^ mice treated with tamoxifen) in response to the serials of Von Frey filaments (**a**) and the brush test (**b**). **c**, Scatterplot of the percentage of paw withdrawal responding to the pinprick test. **d**, Scatterplot of the threshold of tail withdrawal in response to the Randall-Selitto test. **e**, **f**, Scatterplot of the paw withdrawal latencies on the hot plate (**e**) and cold plate (**f**). **g**, Scatterplot of the paw withdrawal threshold in the ramping Von Frey test of baseline (before capsaicin injection) and 15 min post capsaicin injection. **h-j**, Scatterplot of the number of scratches in response to the injected pruritogens including histamine (**h**), chloroquine (**i**) and compound 48/80 (**j**). Data are presented as means ± SEM; the number of mice tested is indicated above each bar. Two-way ANOVA with Bonferroni’s multiple comparisons test for **a**. Unpaired Student’s *t* test for **b**-**j** (WT vs. cKO mice), \**P* < 0.05, \*\**P* < 0.01, \*\*\**P* < 0.001.

We further investigated whether Creld1 is involved in itch sensation. The cKO mice exhibited profound deficits in both histamine-dependent and -independent itch pathways, showing nearly abolished scratching responses to histamine and significantly reduced responses to chloroquine and compound 48/80 (Fig. 3h-j). These results demonstrate Creld1’s essential role in mediating itch perception.

The cKO mice showed normal body weight, motor coordination (rotarod performance), and locomotor activity (open field test) throughout the post-tamoxifen period (Extended Data Fig. 3i-k), confirming that their somatosensory deficits were not secondary to general health or motor impairments. No significant differences were observed in either the proportions of NFH-, IB4-, or CGRP-positive DRG neurons (Extended Data Fig. 3l), or in their projections into the spinal dorsal horn or skin (Extended Data Fig. 3m-p) between WT and cKO mice. Further ultrastructural analysis of the sciatic nerve revealed a mild change of fiber width of myelinated axons in the cKO mice (Extended Data Fig. 3q, r).

Collectively, our comprehensive characterizations of both the sgCreld1-targeted mice and the tamoxifen-induced, Advillin-Cre-mediated conditional knockout mice establishes Creld1 as a master regulator of somatosensation, and also highlight the prowess of the sgRNA-mediated postnatal knockout in overcoming developmental lethality to reveal the somatosensory functions in adult mice.

### Creld1 does not directly modulate the function of sensory receptors

We investigated how Creld1 mediates its broad somatosensory functions. Prior work linked Creld1 to calcineurin/NFATc1 and regulation of ionotropic acetylcholine receptor trafficking^29–31,36^. We examined whether it regulates the expression and function of key sensory receptors. Single-cell calcium imaging revealed no significant difference between WT and cKO DRGs in response to heating, cooling, the TRPV1 agonist capsaicin, the chemical nociceptor TRPA1 agonist AITC, the histamine receptor agonist histamine, the MrgprA3 agonist chloroquine, or 50 mM KCl-induced depolarization (Extended Data Fig. 4a-g). Similarly, whole-cell recordings demonstrated equivalent mechanosensitive current profiles in amplitude and inactivation kinetics (Extended Data Fig. 4h, i). These results suggest Creld1’s sensory regulation occurs independently of direct receptor modulation, pointing instead to potential roles in neuronal excitability, signal integration, or alternative signaling pathways underlying its pan-modal sensory control.

### Creld1 interacts with and regulates voltage-gated sodium channels in mouse DRG neurons

To elucidate Creld1’s molecular mechanism, we performed immunoprecipitation from DRG lysates using the anti-Creld1 antibody followed by mass spectrometry. This analysis specifically identified multiple voltage-gated sodium channel (Na_v_) isoforms (Na_v_1.1, Na_v_1.3, Na_v_1.6, Na_v_1.7, Na_v_1.8, and Na_v_1.9) in the Creld1 pulldown, but not in the IgG control (Extended Data Fig. 5a). Co-immunostaining of Creld1 and Na_v_1.7 showed their co-expression and co-localization in primary DRG neurons, where 73.5 ± 5.1% and 67.2 ± 5.1% of neurons expressed Creld1 and Na_v_1.7 respectively, and 91.8 ± 4.2% of Creld1^+^ neurons co-expressed Na_v_1.7 (Fig. 4a-d, Extended Data Fig. 5g, h). These findings demonstrate a specific interaction between Creld1 and Na_v_ in sensory neurons.

**Fig. 4.**
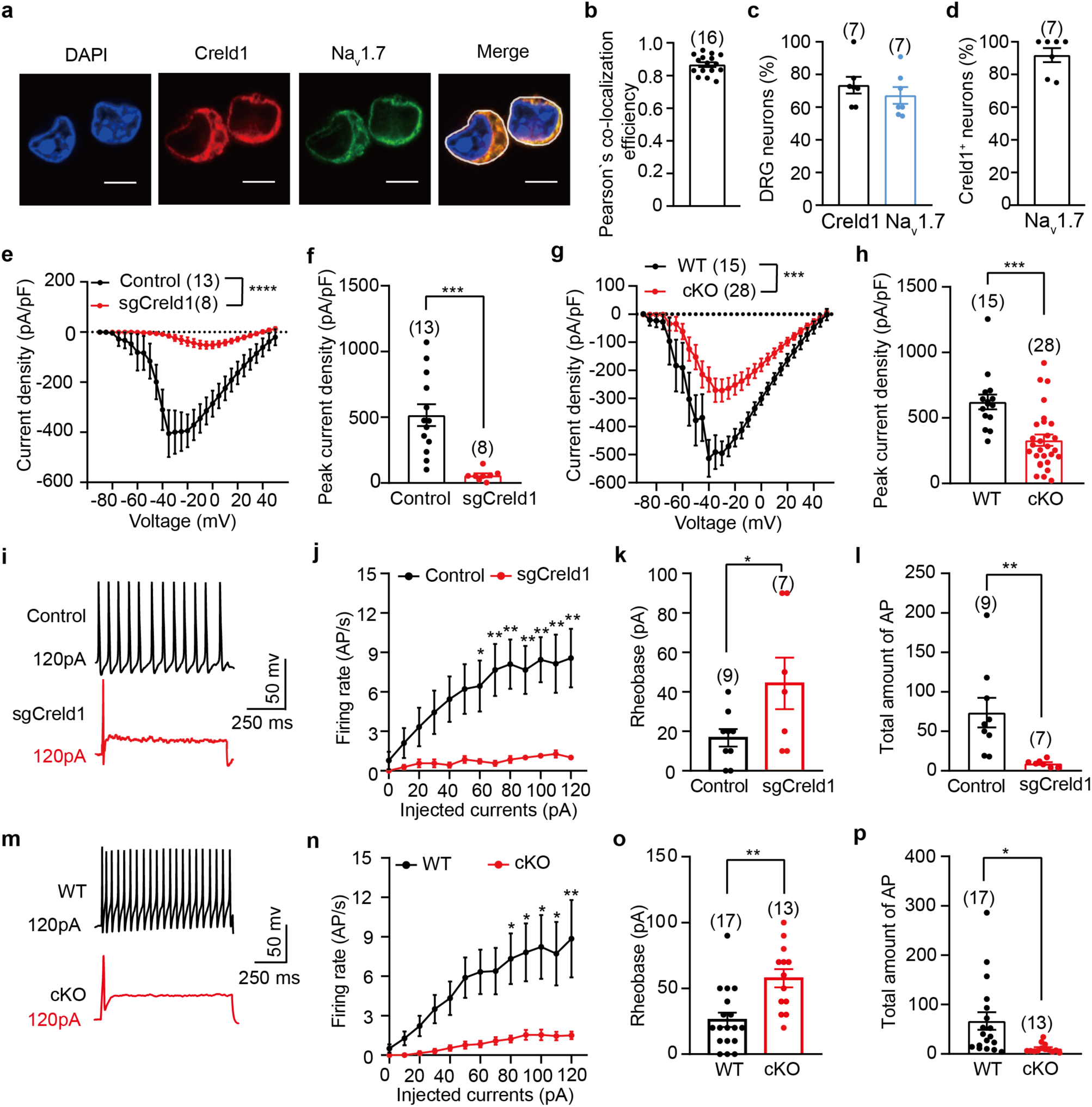
Creld1 regulates Na_v_ and excitability in DRG neurons. **a**, Immunofluorescent staining of endogenous Creld1 and Na_v_1.7 in cultured DRG neurons after 3-12 h in culture using the anti-Creld1 and anti-Na_v_1.7 antibody. Scale bar, 5 μm. **b**, Pearson’s co-localization efficiency analysis for Na_v_1.7 and Creld1 located in DRG neurons. The white circle illustrates the region of interest (ROI) used for analyzing the immunofluorescence staining of Na_v_1.7 and Creld1. Each dot represents an individual cell (16 cells in total)**. c**, Percentage of Creld1-positive or Na_v_1.7-positive DRG neurons (n = 7 sections). **d**, Percentage of Na_v_1.7-positive neurons among Creld1-positive DRG neurons (n = 7 sections). **e**, The I-V curve of control and sgCreld1 DRG neurons. **f**, The peak current density of control and sgCreld1 DRG neurons. **g**, The I-V curve of WT and cKO DRG neurons. **h**, The peak current density of control and sgCreld1 DRG neurons. **i**, Representative trace of evoked APs of control and sgCreld1 DRG neurons. **j**, Firing frequency of control and sgCreld1 DRG neurons across step current injections. **k**, Rheobase of the control and sgCreld1 DRG neurons. **l**, Total amount of APs evoked in control and sgCreld1 DRG neurons. **m**, Representative trace of evoked APs of WT and cKO DRG neurons. **n**, Firing frequency of WT and cKO DRG neurons in response to the step current injections. **o**, Rheobase of WT and cKO DRG neurons. **p**, Total amount of APs evoked in WT and cKO DRG neurons. Data are presented as means ± SEM; the number of cells or sections are indicated. Unpaired Student’s *t* test for **c**, **f**, **h**, **k**, **l**, **o, p**. Two-way ANOVA with Bonferroni’s multiple comparisons test for **e**, **g**, **j**, **n**, \**P* < 0.05, \*\**P* < 0.01, \*\*\**P* < 0.001, \*\*\*\**P* < 0.0001.

To determine whether Creld1 functionally regulates Na_v_, we performed whole-cell recordings in DRG neurons from both control and Creld1-deficient mice. Strikingly, sgCreld1-targeted neurons (indicated by the expression of GFP) exhibited an 89% reduction in peak Na_v_ current density (58.1±14.8 pA/pF) compared to controls (516.2±83.7 pA/pF) (Fig. 4e, f, Extended Data Fig. 5b). This effect was confirmed in DRG neurons derived from the cKO mice (Fig. 4g, h, Extended Data Fig. 5c). Both genetic models demonstrated significantly impaired neuronal excitability, featuring reduced action potential firing rates, elevated rheobase thresholds, and diminished total AP output (Fig. 4i-p). Importantly, qPCR analysis ruled out transcriptional regulation of Na_v_ isoforms (Na_v_1.6-1.9) as the underlying mechanism (Extended Data Fig. 5d). These results establish Creld1 as a crucial regulator of Na_v_ channel function that modulates DRG neuron excitability, thereby explaining its broad influence across multiple somatosensory modalities in mice.

### Structure-function interactions between Creld1 and Na_v_ channels

Creld1 is a membrane protein with a AlphaFold predicted domain organization consisting of a N-terminal signal peptide, a WE domain characteristic of the Creld family, an EGF-like domain, a calcium-binding EGF-like (cbEGF) domain, and two C-terminal transmembrane domains (TM1 and TM2) (Fig. 5a, b). To determine whether it is located on the plasma membrane, we performed cell-surface biotinylation assays and detected Creld1 in the biotinylated fraction by Western blot, confirming plasma membrane localization (Fig. 5c). To resolve its topology, we inserted a Flag tag either at residue F68 (Creld1-F68-Flag), residue D386 (Creld1-D386-Flag) or the C-terminus (Creld1-Flag). The transfected HEK293T cells were subjected to immunostaining of the FLAG tag with or without permeabilizing the cells. Live immunofluorescent staining revealed anti-FLAG immunofluorescent signals on the membrane in Creld1-F68-Flag and Creld1-Flag expressing HEK293T cells, while Creld1 or Creld1-D386-Flag cells showed no signal (Extended Data Fig. 6). Flow cytometry confirmed the topology of Creld1 (Fig. 5d). These results were consistent with an extracellular orientation for both N- and C-termini (Fig. 5a). Collectively, these findings establish Creld1 as a dual-transmembrane protein with extracellular termini, suggesting a potential role in extracellular interactions or signaling.

**Fig. 5.**
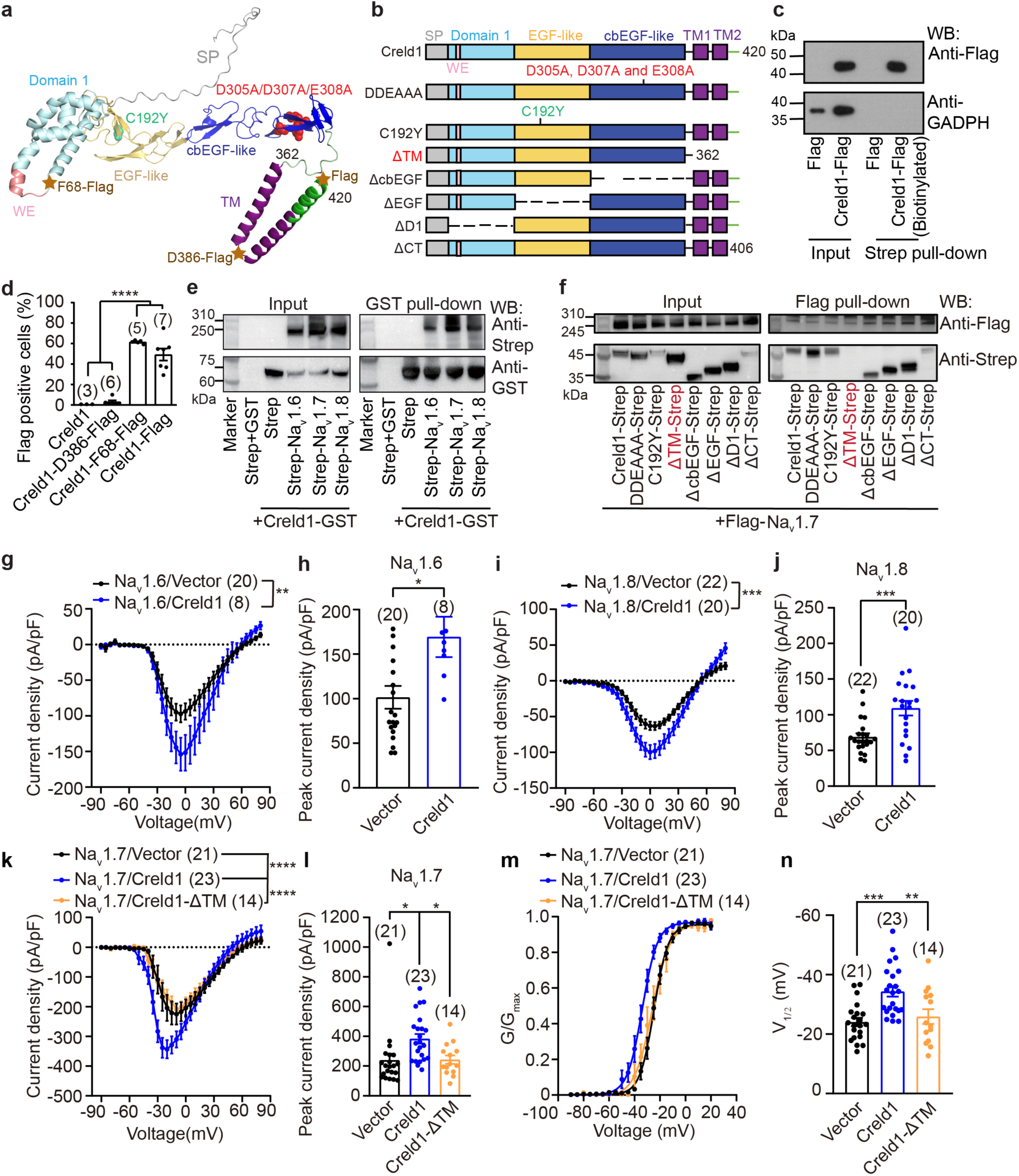
Biochemical and functional characterizations of the Creld1-Na_v_ interaction. **a**, AlphaFold predication of the murine Creld1 protein structure with the indicated domains and the Flag-tag inserted after residue D386 or at the C-terminus labelled with star. **b**, Schematic representation of the indicated functional domains of WT Creld1 and the indicated mutants. **c**, Western blots showing the biotinylated or whole-cell lysate samples from HEK293T cells transfected with the indicated constructs. **d**, Flow cytometry quantification of Flag-positive cells after live staining of constructs with the Flag-tag inserted after residue F68, D386 or at the C-terminus of Creld1-IRES-mCherry. **e**, Western blots showing the interaction between overexpressed Creld1-GST and the indicated isoforms of Na_v_ channels in HEK293T cells. **f**, Western blots showing Flag pull-down of Flag-Na_v_1.7 co-expressed with the indicated Strep-fused Creld1 or mutants. **g**, The I-V curve of HEK293T cells co-transfected with the indicated constructs. **h**, The peak current density of HEK293T cells co-transfected with the indicated constructs. **i**, The I-V curve of D-7/23 cells co-transfected with the indicated constructs. **j**, The peak current density of ND-7/23 cells co-transfected with the indicated constructs. **k**, The I-V curve of HEK293T cells co-transfected with the indicated constructs. **l**, The peak current density of HEK293T cells co-transfected with the indicated constructs. **m**, The voltage dependent activation curves of currents recorded from HEK293T co-transfected with the indicated constructs. **n**, V_1/2_ of the voltage-dependent activation curve of HEK293T cells co-transfected with the indicated constructs. Data are presented as means ± SEM; the number of cells is indicated. One-way ANOVA with Bonferroni’s multiple comparisons test for **d**, **l**, **n**. Unpaired Student’s *t* test for **h**, **j**. Two-way ANOVA with Bonferroni’s multiple comparisons test for **g**, **i**, **k**. \**P* < 0.05, \*\**P* < 0.01, \*\*\**P* < 0.001, \*\*\*\**P* < 0.0001.

We systematically investigated the domain basis of Creld1’s interaction with prominent DRG-expressing Na_v_, including Na_v_1.6, Na_v_1.7 and Na_v_1.8, using heterologous expression in HEK293T cells. GST pull-down assays confirmed binding between Creld1-GST and Strep-tagged Na_v_1.6-1.8 (Fig. 5e), recapitulating our endogenous DRG neuron findings. Through structure-function analysis of various Creld1 mutants, including the disease-associated mutant C192Y, the Ca²⁺-binding-deficient mutant DDEAAA^32^, and the domain-deleted mutants (Fig. 5b), we identified the C-terminal transmembrane (TM) domain as essential for Na_v_1.7 interaction, as deletion of this region containing Met363 to Arg420 (ΔTM) specifically abolished binding while other mutations had no effect (Fig. 5f). These results establish the C-terminal TM domain as the critical interface for Creld1’s regulation of Na_v_ channels, providing mechanistic insight into its control of neuronal excitability.

To determine whether Creld1 influences Na_v_ channel trafficking to the plasma membrane, we performed cell-surface biotinylation assays in HEK293T cells expressing Na_v_1.6, Na_v_1.7, or Na_v_1.8 with or without Creld1. Western blot analysis of streptavidin-purified fractions revealed comparable levels of biotinylated Na_v_ channels regardless of Creld1 co-expression, indicating no effect on surface expression (Extended Data Fig. 7e-j). Interestingly, wild-type Creld1 was efficiently co-purified with biotinylated Na_v_1.6-1.8, while the ΔTM mutant failed to associate with surface Na_v_1.7, confirming the requirement of the C-terminal transmembrane domain for this interaction (Fig. 5e, f). These findings demonstrate that while Creld1 physically associates with Na_v_ channels at the plasma membrane, this interaction appears not to affect their surface trafficking or expression levels.

Electrophysiological characterizations demonstrated that Creld1 significantly enhanced sodium currents mediated by Na_v_1.6, Na_v_1.7 and Na_v_1.8 in heterologous systems, increasing peak current densities by 66.7% for Na_v_1.6, 60.1% for Na_v_1.7, and 58.4% for Na_v_1.8 (Fig. 5g, -l, Extended Data Fig. 7a-c). This functional enhancement required the C-terminal transmembrane domain, as the ΔTM mutant failed to potentiate Na_v_1.7 currents (Fig. 5k, l, Extended Data Fig.7c). Voltage-current analysis revealed that Creld1, but not the Creld1-ΔTM mutant, induced a -10.3 mV hyperpolarizing shift in Na_v_1.7 activation (V_1/2_: -34.4 ± 1.8 mV vs control -24.1 ± 1.4 mV; ΔTM -25.9 ± 2.5 mV) (Fig. 5m, n), demonstrating modulation of channel gating. Collectively, these findings demonstrate that Creld1 biochemically interacts with and functionally regulates Na_v_ channels, and the membrane localization of Creld1 with the TM domain is essential for its regulation of Na_v_.

### Regulation of Na_v_1.7 by human CRELD1 depending on the variable C-terminal TM domain

Human patients carrying mutations in *CRELD1* show neurodevelopmental disorders including early-onset epilepsy, but without reported defects in somatosensory phenotypes^32,37,38^. We therefore asked whether the regulatory effect of mouse Creld1 on Na_v_ is conserved in human CRELD1 (hCRELD1). We aligned the protein sequences of mouse and human Creld1. Notably, hCRELD1 has five isoforms, with the identities to mouse Creld1 ranging from 73.4% to 91.4%: isoform 1 (73.6%), isoform 2 (91.4%), isoform 3 (73.4%), isoform 4 (85.2%), and isoform 5 (76.6%) (Fig. 6a). Among these, isoforms 2 and 4 retain a conserved C-terminal TM domain with that of mCreld1, whereas isoforms 1, 3, and 5 contain highly divergent C-terminal TM domains (Fig. 6a). We cloned hCRELD1 isoform 1 (hCRELD1-Iso1) and isoform2 (hCRELD1-Iso2) for testing their functional and biochemical interaction with Na_v_1.7. Electrophysiological analyses demonstrated that hCRELD1-Iso2 significantly increased Na_v_1.7 current density, mirroring the effect of mCreld1. In contrast, hCRELD1-Iso1 failed to modulate Na_v_1.7 currents compared to controls (Fig. 6b). Furthermore, co-immunoprecipitation experiments revealed an interaction between Na_v_1.7 and hCRELD1-Iso2, whereas no such association was detected with hCRELD1-Iso1 (Fig. 6c, d). Harmonized human somatosensory neuronal cell atlases revealed co-expression of hCRELD1 and Na_v_^27^ (Extended Data Fig. 8). Collectively, these structural-functional analyses demonstrate that human CRELD1 displays isoform-dependent regulation of Na_v_ due to the sequence divergence of the critical C-terminal TM domain.

**Fig. 6.**
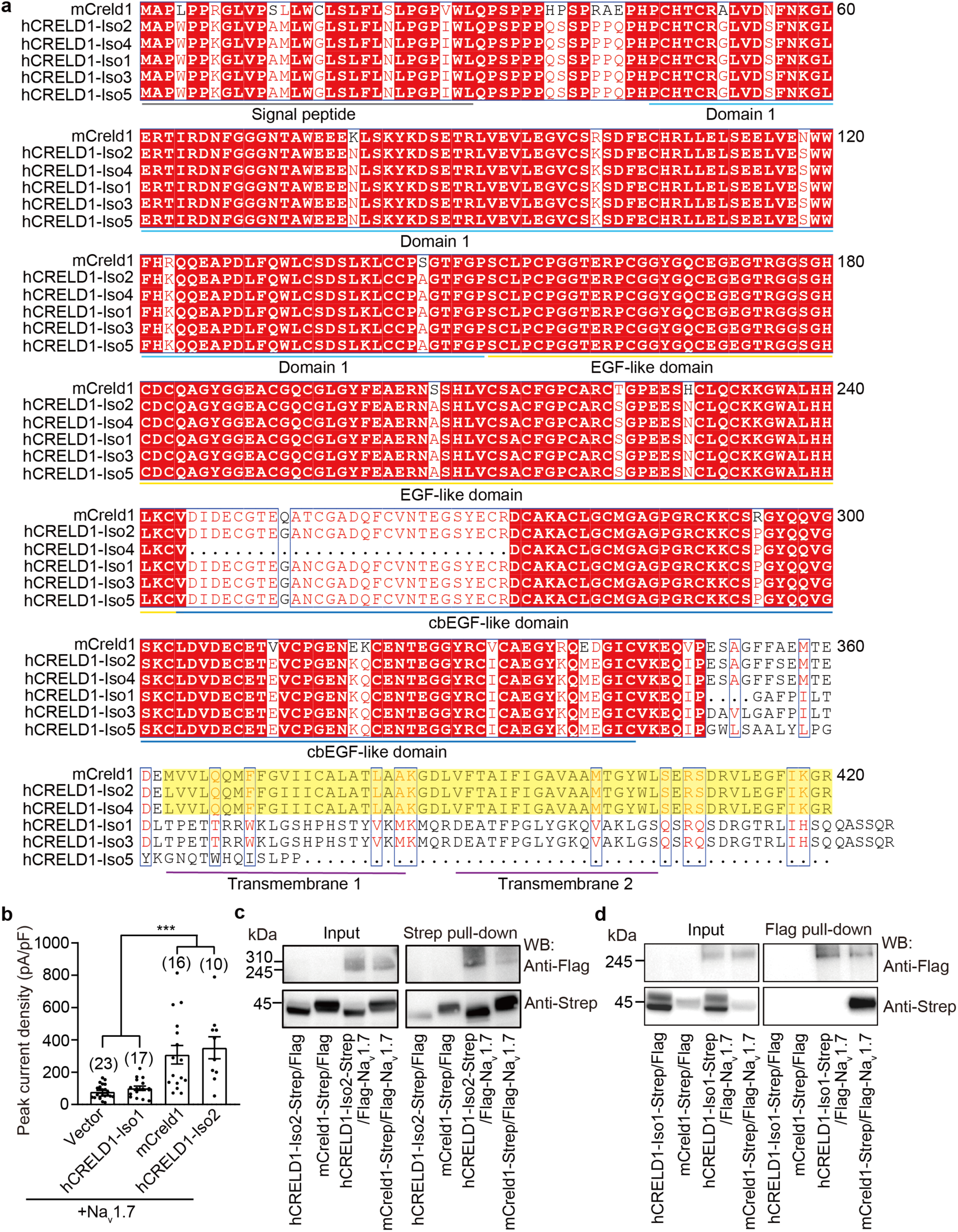
Human Creld1 regulates Na_v_ 1.7 in an isoform-dependent manner. **a**, Sequence alignment of Creld1 derived from mouse and human. The alignment was performed using ClustalW (https://www.genome.jp/tools-bin/clustalw) and Espript 3 (https://espript.ibcp.fr/ESPript/cgi-bin/ESPript.cgi). Notably, human CRELD1 (hCRELD1) have multiple isoforms: isoform 2 and isoform 4 retain a conserved C-terminal TM domain with that of mCreld1 (highlighted in the yellow region), whereas isoform 1, isoform 3, and isoform 5 contain a diversified C-terminal TM domain. Additionally, isoform 4 features a deletion within the cbEGF-like domain when aligned with mouse Creld1. **b**, The peak current density of HEK293T cells co-transfected with the indicated constructs. Data are presented as means ± SEM; the number of cells is indicated. **c**, Western blots showing the co-immunoprecipitation of Flag-Na_v_1.7 by mCreld1-strep and the hCRELD1-Iso2-step. **d**, Western blots showing the co-immunoprecipitation of mCreld1-Strep, but not hCRELD1-Iso1-Strep by Flag-Na_v_1.7. One-way ANOVA with Bonferroni’s multiple comparisons test for **b**, \*\*\**P* < 0.001.

### Overexpression of Creld1 in DRG neurons potentiates somatosensory responses

To determine whether Creld1 can sufficiently modulate endogenous Na_v_ channels in DRG neurons and influence somatosensory behaviors, we intracerebroventricularly injected AAV vectors encoding GFP, Creld1, or the non-interacting Creld1-ΔTM mutant into P0-P2 mice. Whole-cell recordings demonstrated that Creld1-overexpressing neurons exhibited a 35.4% increase in peak sodium current density (743.6 ± 29.4 pA/pF) compared to GFP-expressing controls (555.6 ± 48.2 pA/pF), while Creld1-ΔTM-expressing neurons showed no enhancement (550.8 ± 52.2 pA/pF) (Fig. 7a, b, Extended Data Fig. 7k). Voltage-conductance analysis revealed that Creld1, but not the Creld1-ΔTM mutant, induced a significant -15.1 mV hyperpolarizing shift in the voltage-conductance relationship (V50: -55.5 ± 2.3 mV for Creld1 vs -40.4 ± 3.1 mV for GFP and -42.4 ± 3.4 mV for Creld1-ΔTM) (Fig. 7c, d), mirroring our heterologous expression results with Na_v_1.7 (Fig. 5m, n). These findings demonstrate that Creld1 is both necessary and sufficient to enhance endogenous Na_v_ channel function in sensory neurons through increased current amplitude and altered voltage sensitivity.

**Fig. 7.**
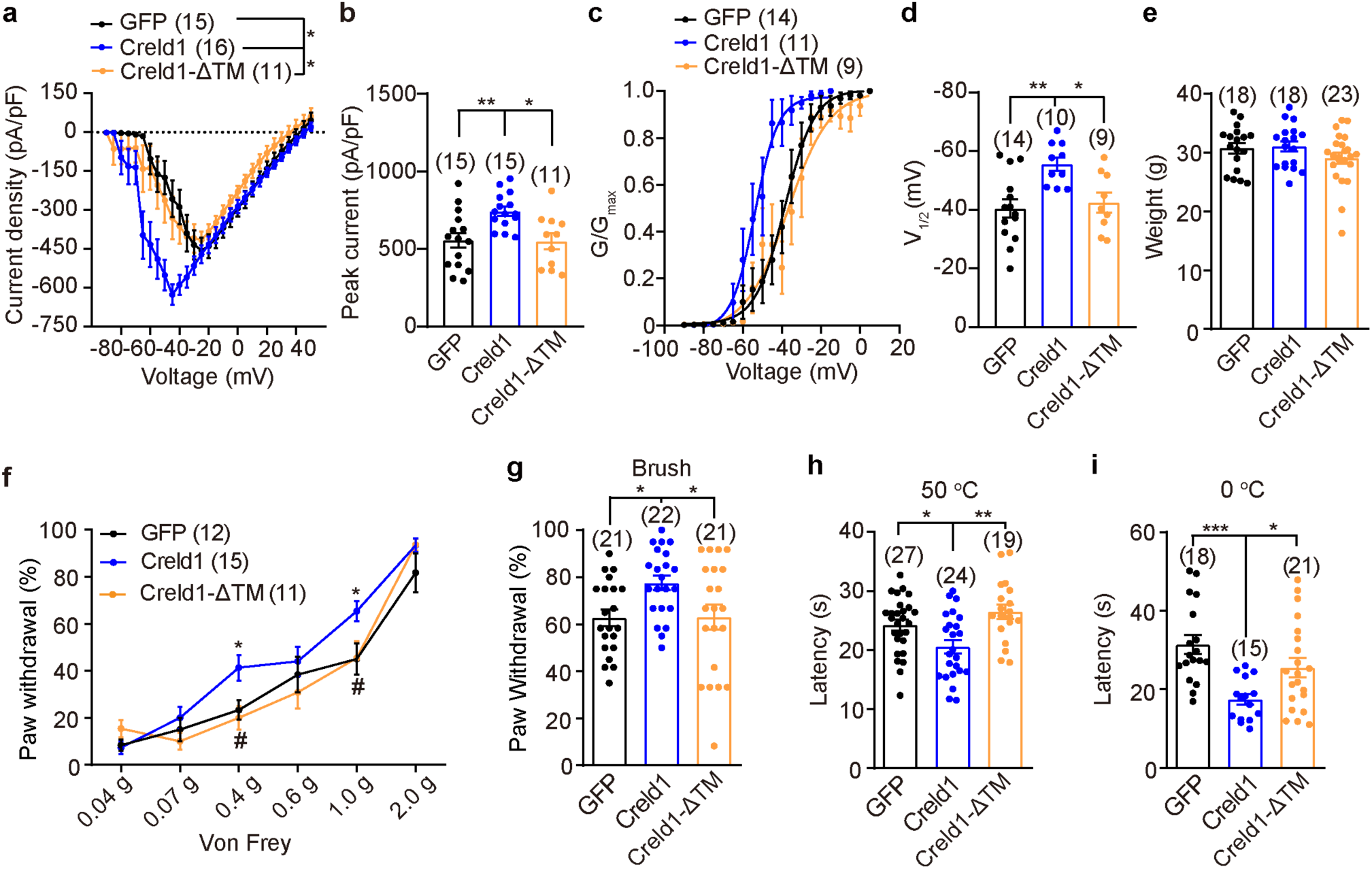
Overexpression of Creld1 enhances somatosensory behaviors. **a**, The I-V curves of Na_v_ currents recorded from DRG neurons derived from the mice injected with AAV9 carrying GFP, Creld1 or Creld1-ΔTM. **b**, The peak current density of Na_v_ currents recorded from DRG neurons derived from the indicated mice. **c**, The voltage-dependent conductance curve of Na_v_ currents recorded from DRG neurons derived from the indicated mice. **d**, V_1/2_ of the voltage-dependent activation curve of Na_v_ currents recorded from DRG neurons derived from the indicated mice. **e**, Body weight of the indicated mice. **f**-**i** Behavioral responses of the indicated mice: Von Frey test (**f**), brush test (**g**), hot plate test (**h**) and cold plate test (**i**). Data are presented as means ± SEM; sample sizes are indicated. Statistical significance is determined by one-way ANOVA with Bonferroni’s multiple comparisons test (**b**, **d**, **e**, **g**, **h**, **i**), two-way ANOVA with Bonferroni’s multiple comparisons test **a**, **f** (*vs. AAV-GFP mice, ^#^vs. AAV-Creld1 mice). \**P* < 0.05, \*\**P* < 0.01, \*\*\**P* < 0.001; ^#^ *P* < 0.05.

Behavioral characterizations of mice overexpressing GFP, Creld1, or Creld1-ΔTM in DRG neurons revealed significant sensory modulation by Creld1. While all groups maintained comparable body weights (Fig. 7e), Creld1-overexpressed mice displayed markedly enhanced responses to Von Frey filaments (Fig. 7f), dynamic brush stimulation (Fig. 7g), and noxious heat or cold stimuli (Fig. 7h, i). Consistent with the increased sodium current density and voltage sensitivity in Creld1-expressing neurons, these behavioral phenotypes demonstrate that Creld1 overexpression potentiates DRG neuronal excitability to amplify responses to light touch and thermal stimuli.

## Discussion

Here we developed an efficient postnatal CRISPR-Cas9 screening platform that overcomes limitations of traditional knockout approaches, enabling systematic functional interrogation of DRG-expressed genes in adult mice. To overcome the potential complication of AAV9 infection in other tissues (Extended Data Fig. 1a), a mouse line with Cas9 specifically expressed in DRG neurons can be selected for the delivery of AAV9-encoded sgRNAs against target genes. Conventional germline deletion of key genes like Piezo2 or Na_v_1.7 leads to perinatal lethality or developmental compensation, hindering the study of their adult sensory functions^2,5^. By contrast, our AAV9-mediated sgRNA delivery achieved acute gene ablation in mature DRG neurons. Remarkably, the sgTrpv1-, sgPiezo2-, and sgNa_v_1.7-targeted mice were not only viable, but also exhibited similar or even more robust behavioral deficits compared to their respective KO mice (e.g. global TRPV1 KO^4,39^, Advillin-Cre-mediated Piezo2 cKO^2^, Na_v_1.8-Cre-, Advillin-Cre or Wnt-Cre-mediated Na_v_1.7 cKO or global KO^5,15,16,20,21^) (Fig. 1). Indeed, the successful identification of Creld1, whose crucial role in adult somatosensation would have remained undiscovered using conventional knockout approaches due to their developmental and perinatal lethality (Fige. 2, 3). Our approach might not totally exclude the possibility of postnatal developmental compensation. However, our observation of postnatal sgRNA-mediated knockout of Trpv1, Piezo2, Na_v_1.7 and Creld1 all generated somatosensory deficits suggest that postnatal developmental compensation appears not to commonly occur. Together, these findings demonstrate the prowess of the postnatal knockout strategy in overcoming the issue of embryonic developmental and perinatal lethality and functional compensation inherently associated with the conventional knockout strategy. Furthermore, given its relatively affordable cost and short time line of experimental execution, this strategy provides a robust framework for high-throughput functional genomics to systematically identify and characterize the full complement of molecular players in sensory neurons, including the long-sought mechanical pain receptors mediating responses to noxious pinch and puncture, which is not mediated by Piezo2 (Ref^40,41^).

Using the postnatal knockout strategy and somatosensory behaviors for a targeted genetic screen, we demonstrated in a proof-of-concept by identifying Creld1 as a master regulator of somatosensation through its modulation of voltage-gated sodium channels in mice, revealing a novel mechanism for tuning neuronal excitability across sensory modalities. Unlike modality-specific receptors (e.g., Trpv1 for heat or Piezo2 for touch), Creld1 knockout broadly impaired responses to mechanical, thermal, pruritic stimuli, and pro-inflammatory substances (Fige. 2, 3). This phenotype mirrors Na_v_1.7 deficiency but contrasts with the selective deficits seen upon Trpv1 or Piezo2 ablation, suggesting Creld1 operates downstream of sensory transductions.

Mechanistically, we found that mouse Creld1 binds Na_v_ channels via its C-terminal TM domain, enhancing current density and hyperpolarizing activation thresholds (Fige. 4–5). Furthermore, mouse Creld1 appears to indifferently bind and regulate different isoforms of Na_v_ channels. The mechanistic regulation of Na_v_ explains its master role in controlling the diverse somatosensory modalities: knockout reduces neuronal excitability (Fig. 4), while overexpression amplifies sensory responses (Fig. 7). Intriguingly, we found that the C-terminal TM domain undergoes a substantial evolutionary divergence from murine to human, resulting in some isoforms of human CRELD1 unable to biochemically interact with and functionally regulate Na_v_1.7 (Fig. 6). Given the uncovered master regulatory role of Creld1 in controlling the somatosenstion of gentle touch, mechanical pain, heat, cold, inflammatory pain and itch in mice (Fig. 2, 3), it is striking that human CRELD1 has evolved to diversify such a robust regulatory mechanism. The reason underlying such an evolutionary selection for human CRELD1 remains unclear. Interestingly, numerous mutations in human CRELD1 have been associated with cardiac and neuronal disorders^31,32,37,38^. Abnormal somatosensory phenotypes of those human patients have not been reported, which could be explained by our finding that some isoforms of human CRELD1 loses its ability to regulate Na_v_1.7. Alternatively, some of the reported mutations might not affect CRELD1’s ability to regulate Na_v_. For instance, the disease-associated C192Y mutation did not alter the biochemical interaction between Creld1 and Na_v_1.7 (Fig. 5f). Nevertheless, it warrants more careful clinical examinations of those human patients for their somatosensory functions.

It has been well known that the expression and biophysical properties of Na_v_ channels are regulated by auxiliary β1 to β4 subunits^42^. Our findings that Creld1 biochemically interacts with and functionally regulate the biophysical properties of Na_v_ channels raises an exciting possibility that Creld1 might function as a novel class of auxiliary subunit. It is worth noting that the hyperpolarizing shift effect of Creld1 on the voltage-dependent activation of Na_v_1.7 or the native Na_v_ channel in DRG neurons is much stronger than the β1 to β4 subunits^43^, indicating differential regulatory mechanisms. Future studies of determining the Creld1-Na_v_ complex structure might provide more detailed understanding of how Creld1 biochemically and functionally regulates Na_v_. Given the long-standing interest of targeting Na_v_1.7 and Na_v_1.8 for developing novel analgesics, the Creld1-Na_v_ interaction interface might offer a precision target for drug development, distinct from blunt Na_v_ channel blockers. Furthermore, our postnatal genetic manipulation of Creld1 expression in DRG neurons opens a novel avenue for gene therapy. Co-delivery of Cas9 and sgCreld1 into specific subtypes of DRG neurons such as nociceptors or itch-mediating neurons might alleviate pain and itch sensation. Beyond somatosensation, Creld1 plays important roles in cardiac development and immunity^29–31,44^. Whether Creld1 modulates Na_v_ function in these systems remains to be addressed as well.

## Methods

### Molecular cloning

The coding DNA for mouse Creld1 (UniProt: Q91XD7) was cloned into the pcDNA3.1 vector using mouse DRG cDNA as the template. Cloning of the human CRELD1 isoform1 and isoform2 was done by Beijing Xianghong Biotechnological Company. The serials of Creld1 mutants were generated. For co-immunoprecipitation and biotinylation assays of cell surface proteins, the Strep tag or GST tag was inserted at the C-terminus of Creld1. For living labelling assays, the Flag tag was inserted after residues D386 or C-terminal end of Creld1. Human Na_v_1.7, human Na_v_1.8, human Na_v_1.9 were gifts from Dr Nieng Yan at Tsinghua University, while human Na_v_1.6 was a gift from Dr Zhuo Huang at Peking University. All constructs, mutations, and truncations of Creld1 were subcloned by using the one step cloning kit under the instruction manual (Vazyme Biotech, C112).

### Animals

All animal experimental procedures were conducted in accordance with the Institutional Animal Care and Use Committee (IACUC) guidelines provided by Tsinghua University. All animals were maintained in a temperature-controlled condition under a 12 h light/dark cycle in the Tsinghua University animal research facility, which is accredited by the Association for Assessment and Accreditation of Laboratory Animal Care International (AAALAC).

CRISPR/Cas9-mediated knockout method was adopted from the previous report^45^. Briefly, Neonatal pups in C57BL/6J background (postnatal days 0-2, P0-P2) of the ROSA26-Cas9-knockin mice (cyagen, China, Stock Number: C001218) were subjected to closed-skull stereotaxic injections. Surgical instruments (27G needles, 10 μL Hamilton syringes) were sterilized with 70% ethanol. Cryoanesthesia was induced by placing pups on crushed ice for 3-4 min, with anesthetic depth confirmed by abolition of toe-pinch reflex. Stereotaxic coordinates (0.7-1.0 mm lateral to sagittal suture, 0.7-1.0 mm caudal to bregma) were marked using sterile surgical ink. Guide RNA (gRNA) was cloned into pAV vector (wzbio, China). The guide RNA sequence for all candidate genes were based on the GeCKO v2 library designed by the Zhang Feng group^43^ and listed (Extended Data Table 1). 1.5 μL of AAV9-sgRNA virus (wzbio, China) was injected at 2 mm depth via microprocessor-controlled microinjection over 60 s. Post-infusion needle retention (20 s) minimized cerebrospinal fluid reflux. Pups recovered on a 37°C warming pad before returning to the home cage.

An embryonic stem (ES) cell targeting construct was designed for generating Creld1^fl/fl^ mice in C57BL/6J background. Two loxP sites flanking exons 3-4 of the Creld1 locus were inserted via homologous recombination in ES cells using a targeting vector with 5’ and 3’ homologous arms (cyagen, China). The Creld1^fl/fl^ mice were crossed with the Piezo2-EGFP-IRES-Cre (the Jaxon Laboratory, Strain: 027719)^2^, Pirt-Cre (a gift from Dr. Xinzhong Dong’s lab)^33^, Advillin-Cre (a gift from Dr. Ardem Patapoutian’s lab)^46^, or Advillin-CreERT2 mice (the Jaxon Laboratory, Strain: 026516, mixed genetic background for less litters when mixed with C57BL/6J inbred mice)^35^ to generate Creld1^fl/fl^ and Advillin-CreERT2/Creld1^fl/fl^ mice. Genotypes were determined by PCR using the primers: Creld1-F, 5’-CATGTCTGGTCAGACTGGGAAG-3’ and Creld1-R, 5’-AATCCGTCATCCCAGCAACTAA-3’ (WT band: 270 bp, fl band: 369 bp); Advillin-CreERT2-F, 5’- CTTTGTGATGTTTCAGTTCCAG-3’ and Advillin-CreERT2-R, 5’-AGGATCTGCACACAGACGGA-3’ (Cre band: 194 bp).

15 mg of tamoxifen (Sigma, T5648) was dissolved into 80% corn oil: 20% ethyl alcohol and made freshly before use. Creld1^fl/fl^ and Advillin-CreERT2/Creld1^fl/fl^ mice of six weeks were both intraperitoneally injected with tamoxifen at 150 mg/kg every other day for three times. Somatosensory behavioral assays were performed on mice between 14 and 35 days after tamoxifen injections.

AAV9-*hsyn*-GFP (AAV-GFP, 4.93 x 10^13^ virions/mL), AAV9-*hsyn*-mCreld1 (AAV9-Creld1, 8.21 x 10^13^ virions/mL, UniProt: Q91XD7) and AAV9-hsyn-Creld1-ΔTM (AAV-Creld1-ΔTM, 8.17×10^13^ virions/mL) were manufactured in the WZ Biosciences Inc company (wzbio, China). 1.5 μL of AAV9 virus was injected into each ventricle of neonatal pups (P0-P2) as the above description of CRISPR/Cas9-mediated knockout method. Mice at 8-16 weeks were used for behavioral assays.

### Primary DRG neuron culture

DRGs were harvested from adult mice (4-8 weeks) euthanized via avertin (250 mg/kg, intraperitoneal injection). Ganglia were dissected within 30 min post-mortem in ice-cold Dulbecco’s Modified Eagle Medium/Nutrient Mixture F-12 (DMEM/F-12 medium, ThermoFisher Scientific, C11330500BT). The dissected DRGs were sequentially incubated in 1.25% collagenase IV (ThermoFisher Scientific, 17104019) for 45 min, then 1.25 unit/ml papain (Worthington, LS003126) for 30 min. Tissues were mechanically dissociated via 1 ml pipettes in DMEM/F-12 medium supplemented with 10% horse serum (Solarbio, S9050) and 100 U/mL penicillin-streptomycin (ThermoFisher Scientific, 15140-122). Cell suspensions were purified through a 15% bovine serum albumin (BSA, Sigma A4503) density gradient. Neurons were plated on poly-D-lysine (0.1 mg/mL, Beyotime, C0312)/laminin (5 μg/mL, Sigma, L2020)-coated coverslips in defined culture medium: DMEM/F12 supplemented with 10% horse serum, 100 U/mL penicillin-streptomycin, 100 ng/mL β-nerve growth factor (NGF, ThermoFisher Scientific, 13257019), and 5 ng/mL glial-derived neurotrophic factor (GDNF, ThermoFisher Scientific PHC7044). Cultures were maintained at 37°C in 5% CO₂ humidified incubator for 3-12 h prior for action potential recording and 12-36 h prior to other functional assays.

### Fura-2 single cell Ca^2+^ imaging

Primary cultured DRG neurons grown on 8 mm coverslips were subjected to single-cell Ca^2+^ imaging^47^. In brief, cells were washed with the Ca^2+^ imaging buffer (1 x HBSS buffer with 1.3 mM Ca^2+^, 10 mM HEPES, pH 7.2) and then loaded with the Ca^2+^ imaging buffer containing 2.5 mM Ca^2+^ indicator dye Fura-2, AM (ThermoFisher scientific, F1225) and 0.05% Pluronic F-127 (Beyotime, ST501). 30 min after Fura-2 loading, cells were washed twice with the Ca^2+^ imaging buffer. The Fura-2 signals excited at either 340 nm or 380 nm were captured by an inverted Nikon-Tie microscope with a CCD camera and DG-4 light box (Sutter Instrument). Intracellular Ca^2+^ concentration was indicated as the 340/380 ratio by using MetaFluor Fluorescence Ratio Imaging software (Molecular Device). To measure TRPV1 function in control and sgTRPV1 DRG neurons, 10 μM capsaicin was applied in Fig. 1b, c. To calculate temperature responses for WT and cKO DRG neurons, the temperature change of bath solution was controlled using a CL-100 temperature controller (Warner Instruments) and a SC-20 Solution In-Line Heater/Cooler (Harvard Apparatus), and recorded with a thermistor placed at the outlet of the perfusion. Various compounds, including 10 μM capsaicin, 100 μM AITC, 50 μM histamine, 1 mM chloroquine and 50 mM KCl were used to calculate the function of WT and cKO DRG neurons.

### Immunohistochemistry

Adult mice were anesthetized by intraperitoneal injection of avertin. DRGs or other tissues were dissected from mice then fixed in 4% PFA for 2-8 hours (h) at 4 °C, and incubated overnight in 30% sucrose at 4 °C. The processed tissues were then embedded in Tissue-Tek O.C.T compound and cryo-sectioned into 5-10 μm sections (DRGs and vagus ganglion) or 30-50 μm sections (brain, spinal cord and footpad skin). Before staining, sections were dried in 60 °C for 20 min, then washed with phosphate-buffered saline (PBS) containing 0.2% Triton X-100 for 30 min, and blocked in PBS with 3% BSA for 30 min. The sections were incubated with primary antibody (diluted in 3% BSA or 1% BSA/5% donkey serum) at 4 °C for overnight, washed with PBS containing 0.1% Tween-20 (PBST) three times, and subsequently incubated with secondary antibodies for 1 h at room temperature. After washed with PBST three times, the sections were incubated with DAPI, dried and mounted with anti-fade mounting medium (ThermoFisher Scientific, P36980). Primary antibodies: rabbit anti-GFP (ThermoFisher Scientific, A11122, 1:500 dilution), chicken anti-GFP (Aves Labs, GFP-1020, 1:500 dilution), chicken anti-NFH (Aves Labs, NFH, 1:500 dilution), biotin IB4 (ThermoFisher Scientific, I21414, 1:100 dilution), rabbit anti-CGRP (immunostar, 24112, 1:500 dilution), mouse anti-Piezo2 (generated in our lab, 1:500 dilution), mouse anti-Trpv1 (Abcam, ab203103,1:500 dilution), human anti-Na_v_1.7 (US11643458B2, a gift from Dr. Juanjuan Du at Tsinghua university, 1:500 dilution), goat anti-Creld1 (R&D systems, AF4116, 1:500 dilution), rabbit anti-TUBB3 (CST, 5666S, 1:500 dilution), rabbit anti-PGP9.5 (Cell signaling technology, 13179). Secondary antibody: AlexaFluor-488 donkey anti-goat secondary antibody (ThermoFisher Scientific, A-11055), AlexaFluor-488 donkey anti-rabbit secondary antibody (ThermoFisher Scientific, A-21206), AlexaFluor-488 donkey anti-chicken secondary antibody (Jackson lab, 703-545-155), Goat Streptavidin Alexa Fluor® 488 secondary antibody (Jackson lab, 016-540-084), AlexaFluor-488 goat anti-human (ThermoFisher Scientific, A11013), AlexaFluor-568 donkey anti-rabbit (ThermoFisher Scientific, A10042), Cys3 donkey anti-chicken (Jackson lab, 703-165-155), Streptavidin Alexa Fluor 594 (Jackson lab, 016-580-084), AlexaFluor-568 goat anti-human (ThermoFisher Scientific, A21090), AlexaFluor-568 goat anti-mouse (ThermoFisher Scientific, A11004), AlexaFluor-647 donkey anti-goat secondary antibody (ThermoFisher Scientific, A-21447), all secondary antibodies were diluted to 1:500. These procedures are also suitable for cultured DRG neurons or permeabilized HEK293T cells transfected with D386-Flag-Creld1 or Creld1-Flag plasmids. All the imaging procedures were performed on the Nikon A1 HD25 confocal microscope. The images were analyzed using the Nikon NIS-Elements AR software or ImageJ software.

### Hybridization chain reaction (HCR) staining

For each target mRNA, a kit containing a DNA probe set, a DNA HCR amplifier, and hybridization, wash and amplification buffers was purchased from Molecular Instruments (moleculartinstruments.com), a non-profit academic resource within the Beckman Institute at Caltech. Sequences for HCR amplifiers B1 is given in the literature^48,49^. Frozen tissue sections were processed for dehydration using ethanol according to the HCR RNA-FISH protocol provided by Molecular Instruments. For probe hybridization, samples were pre-incubated in Probe Hybridization Buffer at 37 °C for 10 minutes, followed by overnight incubation with 0.4 pmol/μL Piezo2 or Trpv1 probes (Molecular Instruments) in 100 μL of Probe Hybridization Buffer. Samples were then washed using Wash Buffer and 5× SSC containing 0.1% Tween-20. For the amplification stage, samples were pre-incubated at room temperature in Amplification Buffer for 30 minutes. Separately, 6 pmol each of hairpin h1 and hairpin h2 (Alexa 647) were prepared by snap cooling: 2 μL of 3 μM stock solution was heated at 95 °C for 90 seconds, then cooled to room temperature in the dark for 30 minutes. The prepared hairpins were then added to 100 μL of Amplification Buffer, and the samples were incubated for 6 hours at room temperature. Images were taken on a A Zeiss LSM 710 inverted confocal microscope.

### Cell culture and transfection

For co-immunoprecipitation between Creld1 and Na_v_ channels, HEK293T cells were cultured in Dulbecco’s Modified Eagle Medium (DMEM, ThermoFisher scientific, C11995500BT) supplemented with 10% (vol/vol) fetal bovine serum (ThermoFisher Scientific) and 100 U/ml penicillin-streptomycin (ThermoFisher Scientific, 15140122) in 15 cm^2^ dishes. HEK293T cells were transfected using polyethylenimine (PEI) (MCE, HY-K2014). After reaching approximately 90% confluence, cells were transfected with plasmids. For co-transfection of Creld1 and Na_v_1.6-Na_v_1.8 plasmids, a total of 30 µg plasmids DNA (1:1 mass ratio) and 96 µL PEI were separately diluted in 1.5 mL Opti-MEM (ThermoFisher Scientific, 51985034), gently mixed, and incubated for 5 min at room temperature. The diluted plasmid and PEI solutions were then combined, mixed gently, and incubated for further 20 min. The DNA/PEI mixtures were added dropwise to the cells. Transfected cells were harvested 36 h post-transfection.

HEK293T cells and ND-7/23 cells were used for electrophysiology of Na_v_1.6-Na_v_1.8. HEK293T cells were cultured in DMEM supplemented with 10% (vol/vol) fetal bovine serum and 100 U/ml penicillin-streptomycin while ND-7/23 cells were cultured in DMEM medium supplemented with 4.5 mg/mL glucose and 10% (vol/vol) fetal bovine serum. Cells were plated onto 8-mm glass poly-d-lysine-coated coverslips in 24-well plate and transfected with 1 µg Creld1/Na_v_ plasmids mixture (1:4 mass ratio). Cells transfected with virous plasmids were sub-cultured after 24 h and examined using electrophysiology.

### Co-immunoprecipitation (Co-IP)

Cells were washed twice with PBS containing 2 mM EDTA (Solarbio, E1171), scraped, and spin at 3,000 × g for 8 min at 4 °C. Pellets were washed twice and lysed in 1 mL lysis buffer containing 25 mM NaPIPES, 140 mM NaCl, 1.0 mM EGTA, 1% CHAPS, 0.5% phosphatidylcholine, 0.05% C12E9, 2.5 mM dithiothreitol (DTT), 1 x protease inhibitor (PI, Roche, 04693132001). Samples were gently rotated at 4 °C for 2 h. Beads (glutathione magnetic agarose, ThermoFisher scientific, 88816; streptavidin magnetic beads, ThermoFisher scientific, 78601; Anti-Flag magnetic beads, Beyotime, P2115) were pre- equilibrated in lysis buffer after washing twice in PBS with 0.5% Tween-20 (Biobying, 2022Y8745). Lysates were centrifuged at 12,000 × g for 20 min at 4 °C, and 10% of the supernatant was retained as the input. The remainder was incubated with pre-washed beads overnight at 4 °C. Beads were washed three times with wash buffer containing 25 mM NaPIPES, 140 mM NaCl, 0.05% C12E9, 0.01% phosphatidylcholine, 2.5 mM DTT, 1 x PI, and bound proteins were eluted in 50 µL 1× SDS loading buffer (Epizyme, LT101S) at room temperature for 30 min. Input samples were denatured similarly.

### Protein extraction from tissue

DRGs were dissected from mice then washed twice with PBS, and rapidly frozen in liquid nitrogen. Frozen tissues were placed in a pre-cooled mortar, ground to powders using a pestle, and transferred to 1.5 ml centrifuge tubes. The tissue powder was lysed in 150-200 µL of RIPA lysis buffer (Beyotime, P0038) supplemented with 1 x protease inhibitor (PI, Beyotime, P1045) by vortex, followed by incubation at 4°C for 30 min. Lysates were clarified by centrifugation at 12,000 × g for 30 min at 4°C. The supernatant was collected; a portion was used for protein concentration determination via bicinchoninic acid (BCA) assay kit (ThermoFisher Scientific, 23227), and the left portion was mixed with 5 × protein loading buffer (Epizyme, LT101S) and denatured at 100°C for 5-10 min.

### Biotinylation of cell surface proteins

Cells were washed with 1 x PBS containing 0.49 mM MgCl₂ and 0.9 mM CaCl₂. Then cells were incubated with 0.5 mg/ml Sulfo-NHS-LC-Biotin (ThermoFisher Scientific, 21217) at 4 °C in the dark for 45 min. Excess NHS-biotin was quenched with 100 mM glycine for 5 min. Cells were washed and lysed. 10% of the lysates were used as the whole-cell lysate samples and the remaining cell lysates were incubated with streptavidin magnetic beads (ThermoFisher scientific, 78601) at 4 °C overnight. After 5 times of washing, the precipitated sample was denatured and prepared for SDS-PAGE gel separation and western blotting.

### Western blotting

Equal amounts of protein (20 µL) were separated by 4–12% SDS–PAGE gradient gels (Epizyme, LK307) at 120 V for 2 h and transferred to PVDF membranes using a rapid transfer buffer (MF649-01) under constant current (400 mA) for 40 min. Membranes were blocked with 5% (w/v) skim milk in Tris buffered saline with 0.1%Tween-20 (TBST) for 1 h at room temperature and incubated overnight at 4 °C with primary antibodies diluted in blocking buffer. Primary antibodies included anti-Flag (Vazyme, 7E701C1,1:3000 dilution), anti-GST (CST, 2622S, 1:10000 dilution), anti-Strep (EASYBIO, BE2076, 1:3000 dilution), anti-GAPDH (Yeasen, 334078,1:3000 dilution), anti-HSP70 (Yeasen, T55496M, 1:3000 dilution), anti-Creld1(R&D systems, AF4116, 1:500 dilution), anti-Na_v_1.7 (a gift from Prof. Juanjuan Du, Tsinghua University, 1:500 dilution) and anti-actin (CST, 7074S, 1:3000 dilution). After three times 10-min TBST washes, membranes were incubated with HRP-conjugated secondary antibodies (anti-mouse IgG, CST, 7076 1:20000; anti-rabbit IgG, CST, 7074S, 1:10,000 dilution; anti-goat IgG, Yeasen, 80790730,1:3000 dilution) at room temperature for 1 h. Signals were visualized using enhanced chemiluminescence (ECL) substrate (Epizyme, SQ201) and detected by Tanon 5200 automatic luminescence imaging system and analyzed by ImageJ software.

### RNA extraction and quantitative real-time PCR (qRT-PCR)

Mouse heart, brain, DRG, skin and Neuro-2a (N2A) cells were collected, flash-frozen in liquid nitrogen and ground to powder. Total RNA was isolated using the RNA extraction kit (Uelandy, UE-MN-MS-RNA-50) according to the manufacturer’s protocol. Total RNA (2 μg) was reverse-transcribed using the HiScript III All-in-One RT SuperMix kit (Vazyme, R333-01) following the manufacturer’s instructions. Primers for all qPCR experiments were listed (Extended Data Table 2). qPCR was performed using the SYBR Green Pro Taq HS Premix qPCR kit III (low Rox; AG11739) in a 20 µl reaction volume. Amplification was conducted according to the manufacturer’s protocol. Briefly, cycling conditions were as follows: Hold on 50 °C for 2 min and 95 °C for 10 min, followed by 40 cycles of 15 s at 95 °C and 60 s at 60 °C. Melting curve analysis was performed at 95 °C for 15 s and 65 °C for 15 s. Expression of Creld1, Na_v_1.6, Na_v_1.7, Na_v_1.8, Trpv1, or Piezo2 was normalized to β-actin. The results were analyzed by the ViiA™ 7 Real-Time PCR software, and the gene expression was determined using the comparative Ct method^50^. The mRNA levels were expressed as normalized mRNA levels.

### Mass spectrometry

DRGs were collected from two mice of 8-16 weeks and rapidly frozen in liquid nitrogen. Frozen tissues were placed in a pre-chilled mortar, ground to a powder using a pestle, and transferred into lysis buffer containing 25 mM NaPIPES, 140 mM NaCl, 1.0 mM EGTA, 1% CHAPS, 0.5% phosphatidylcholine, 0.05% C12E9, 2.5 mM dithiothreitol (DTT), 1 x Protease inhibitor. 0.25 mg protein A/G magnetic beads (ThermoFisher scientific, 88803) were incubated with either 10 µg normal goat IgG or 10 µg anti-Creld1 antibody at 4 °C for 2 h. Then the antibody-bound beads were incubated with DRG cell lysates at 4 °C overnight. After incubation, the beads were washed for 5 times with the wash buffer containing 25 mM NaPIPES, 140 mM NaCl, 0.05% C12E9, 0.01% phosphatidylcholine, 1 x PI. The protein samples were eluted with 50 µL 1 x protein loading buffer, separated on the 12% SDS-PAGE gel and visualized by silver staining kit (Sigma-Aldrich). Gel lanes corresponding to each sample were excised and subjected to mass spectrometry identification in the Protein Core Facility of Tsinghua University.

### Transmission electron microscope

Sciatic nerves from WT (n=3) and cKO (n=3) mice were isolated and fixed in 2% paraformaldehyde/2.5% glutaraldehyde in 0.1 M PBS at room temperature for 1 h. The fixed samples remained on ice were sent to Cell Biology Facility, center of Biomedical Analysis of Tsinghua University, examined by a Hitachi (HT) 7800 transmission electron microscope and analyzed by Dragonfly and Arivis.

### Electrophysiology

Patch-clamp experiments were performed using HEKA EPC10 as previously described^1,11,51–54^. Approximately equal numbers of small, medium and large size of DRGs were chosen for recording mechanically-induced currents, Na_v_ currents and action potential. All experiments were done at room temperature. The recording electrodes using in all experiments had a resistance of 3-5 MΩ. Currents were sampled at 20 kHz for whole-cell mechanosensitive currents and current-clamp recording and 50 kHz for whole-cell Na_v_ currents recording, filtered at 3 kHz using PatchMaster software. Leak currents were subtracted off-line from the current traces. Data were analyzed using GraphPad Prism and Fitmaster software.

For whole-cell mechanosensitive currents recordings in DRG neurons, the standard extracellular solution was composed of (in mM) 133 NaCl, 3 KCl, 2.5 CaCl_2_, 1 MgCl_2_, 10 glucose and 10 HEPES (pH 7.3 with NaOH). The internal solution composed of (in mM) 133 CsCl, 1 CaCl_2_, 1 MgCl_2_, 5 EGTA, 4 MgATP, 0.4 Na_2_GTP and 10 HEPES (pH 7.3 with CsOH). Mechanical stimulation to DRGs was delivered at an angle of 80° using a fire polished glass pipette (tip diameter 3 - 4 mm) as described^1,11,51,52^.

For Na_v_ currents and tetrodotoxin (TTX)-S currents recordings in DRG neurons, the standard extracellular solution was composed of (in mM) 150 NaCl, 2 KCl, 1.5 CaCl_2_, 1 MgCl_2_, 10 glucose and 10 HEPES (pH 7.4 with NaOH). The internal solution composed of (in mM) 135 CsF, 10 NaCl, 1 EGTA and 5 HEPES (pH 7.4 with CsOH). 1 µM TTX was used in the bath solution to isolate the TTX-S currents. The TTX-S currents were obtained by the subtraction of the sodium currents obtained before and after TTX treatment.

Na_v_1.6, Na_v_1.7 and its variants were recorded in HEK293T cells, Na_v_1.8 was recorded in ND-7/23 cells due to its small peak currents in HEK293T cells. 400 nm TTX was added to the bath solution during recording of ND-7/23 cells to completely block endogenous TTX-sensitive sodium current. The standard extracellular solution was composed of (in mM) 140 NaCl, 4 KCl, 1.5 CaCl_2_, 1 MgCl_2_, 10 D-glucose and 10 HEPES (pH 7.4 with NaOH). The internal solution composed of (in mM) 105 CsF, 40 CsCl, 10 NaCl, 10 EGTA and 10 HEPES (pH 7.4 with CsOH).

In both HEK293T cells transfected with Na_v_1.6 or Na_v_1.7 and ND-7/23 cells transfected with Na_v_1.8, the voltage dependence of ion current (I-V) was analyzed using a protocol consisting of steps from a holding potential of -100 mV to voltages ranging from -90 to +80 mV (+50 mV for DRG neurons) for 100 ms in 5-mV increments. The linear component of leaky currents and capacitive transients were subtracted using the P/4 procedure. Activation curves were obtained by calculating conductance (G) at each voltage (V) using the equation G = I/(V – V_r_), where V_r_ represents the reversal potential (the voltage at which the current is zero). For activation curves, conductance was normalized and plotted against the maximal sodium conductance. Activation curves were fitted with the Boltzmann function to obtain V_1/2_ and slope values.

For current-clamp recording, pipettes were filled with the internal solution contained (in mM): 136 K-Glucose,10 NaCl, 1 MgCl_2_, 10 EGTA, 10 HEPES and 2 Mg-ATP, adjusted to pH 7.3 with KOH. The extracellular solution contained (in mM): 154 NaCl, 5.6 KCl, 1 MgCl_2_, 2 CaCl_2_, 10 Glucose and 8 HEPES, adjusted to pH 7.4 with NaOH. A whole-cell configuration was obtained at room temperature in voltage-clamp mode and then the recording was performed after switching to current-clamp mode. Action potentials (APs) were induced by 1 s somatic current injections starting from 0 pA to 120 pA in 10-pA increments. Rheobase was selected as the first current step which was sufficient to initiate APs.

### Live cell labeling

HEK293T cells were cultured on 8 mm coverslips and transfected with 1 µg Creld1, Creld1-F68-Flag, Creld1-D386-Flag or Creld1-Flag plasmids. 36 h later, cells were incubated with the mouse anti-FLAG antibody (Abmart, M20008, 1:100 dilution) at 37°C for 20 min or room temperature for 1 h, treated with the AlexaFluor-488 donkey anti-mouse IgG secondary antibody (ThermoFisher Scientific, A-21202, 1:200 dilution) at room temperature for 1 h and fixed with 4% paraformaldehyde (PFA). For cell permeable staining, cells were fixed with 4% PFA, permeabilized with 0.02% Triton X-100 and blocked with 3% bovine serum albumin (BSA). Following incubation with the mouse anti-Flag antibody (Abmart, M20008, 1:500 dilution) and the AlexaFluor-488 donkey anti-mouse IgG secondary antibody (ThermoFisher Scientific, A-21202, 1:500 dilution), coverslips were mounted onto the glass slides. Images were acquired using Nikon A1 confocal microscopy.

### Flow cytometry

HEK293T cells were cultured on 24-well-plate and transfected with 1 µg Creld1, Creld1-F68-Flag, Creld1-D386-Flag or Creld1-Flag plasmids. 36 h later, cells were washed and incubated with the mouse anti-FLAG antibody (Abmart, M20008, 1:100 dilution) at 37°C for 20 min or room temperature for 1 h, washed and treated with the AlexaFluor-488 donkey anti-mouse IgG secondary antibody (ThermoFisher Scientific, A-21202, 1:200 dilution) at room temperature for 1 h. Samples were resuspended in PBS containing 2% FBS, filtered through a 70-µm filterand analyzed using BD FACSAriaII (BD Biosciences).

### Von Frey filament and brush tests

Male mice (8-16 weeks) were acclimated in transparent Plexiglas chambers (Ugo Basile 46000) on an elevated wire grid (5 mm mesh) for 30 min/day over two consecutive days. Mechanical hypersensitivity was assessed using a graded series of Von Frey filaments (0.008 - 2.0 g; Bioseb BIO-VF-M) applied perpendicularly to the plantar hind paw through the mesh floor. Each filament strength was tested six times with 30 s intervals, following the up-down method^55^. Withdrawal frequency (%) was calculated as (number of positive responses/6 trials) × 100. Similarly, soft paint brush was gently applied to the plantar of hind paw, and withdraw percentage out of six trials was counted. The investigator was blinded to the genotypes throughout testing.

### Ramping Von Frey

Mice were acclimated and tested using the transparent Plexiglas chambers (Ugo Basile). A ramping protocol was applied that increased in force gradually from 0 - 25 g over the course of 10 s. The filament was applied to the plantar surface of the hind paw until a withdrawal response was observed. Each trial was repeated three times. Baseline measurements were first recorded for all mice. Subsequently, mice received a hind paw injection of 20 μL Complete Freund’s Adjuvant (CFA; Sigma, F5881) and were retested 24 h post-injection. In a separate experiment, the mice were treated with 10 μL of 0.5 mM capsaicin (Sigma, 211275) in the hind paw and tested 15 min after administration. For each mouse, the three force readings were averaged, and the mean withdrawal threshold was plotted for analysis.

### Randall-Selitto assay

Mechanical noxious threshold was determined by Randall-Selitto device (Ugo Basile, Product ID37215). Briefly, mice were accumulated in a plastic tube for 10 min two days before testing. A The maximum pressure was set to 250 g for preventing tail damage. Slowly increasing pressure was applied to the tail by the blunt tip of the device until the mice exhibited tail flicking or escape behavior. The threshold was recorded. Four trails were carried out with 5-10 min intervals and the threshold was calculated by averaging the four trails.

### Pinprick test

Following two-day habituation (30 min/day), A 2ml syringe needle was applied to the glabrous skin of the hind paw, taking care not to pierce through the skin. Each mouse was tested ten times with interval of 1 min. Paw withdrawal, shaking, or licking was scored as a positive response and reported as percentage for the total number of trials.

### Hot plate test

Mice were placed on a 50.0 ± 0.2°C aluminum plate (Bioseb BIO-HP-LE) enclosed by a clear acrylic cylinder (20 cm diameter). Latency of hind paw licking and jumping was recorded with 60 s cutoff.

### Cold plate test

The cold plate was set to 0 °C and -5.0°C (Bioseb BIO-CO-LE), and the latency of fore paw lifting, hind paw licking or jumping was recorded within 60 s maximum exposure.

### Itch-evoked scratching behavior

Two days prior to experimentation, mice were anesthetized with 1–2% isoflurane and shaved on the nape of the neck and right cheek using surgical clippers. On two consecutive days before testing, mice were acclimated individually for 30 min in covered four-chamber plexiglass arenas with opaque dividers (Ugo Basile). To evoke scratching behavior, mice received subcutaneous injections of either: (1) Chloroquine (200 μg in 50 μL PBS; Sigma) or Compound 48/80 (100 μg in 50 μL PBS; Sigma) in the neck, or (2) Histamine (50 μg in 10 μL PBS; Sigma) in the cheek. A bout of scratching was defined as a continuous episode of hind paw movement directed toward and away from the injection site. All behaviors were video-recorded for 30 min, and scratching bouts were quantified.

### Rotarod test

Mice were trained on the rotarod device (KEW Basis, China, product ID KW-600X) following a protocol of staying on the rotarod at 5 rpm for entail 5 min for two days as the previous described^52^. After two days training, the rotarod started at 5 rpm and accelerated to 40 rpm over 5 min. The latency of the mice falling of the rotarod was automatically recorded. Each mouse was tested three times with interval of 30 min, and the average was considered as the rotarod latency.

### Open field test

The open field test was conducted using an acrylic box (50 × 50 × 40 cm). Mice were acclimated to the behavioral testing room for 30 min, transferred to the acrylic box (one mouse per box) and followed by a 10-min video-recording of their unrestricted movement. Subsequently, the videos were analyzed using Noldus EthoVision XT software to quantify the total distance traveled over the 10-min period, as well as the duration of time spent in the central zone.

### Statistical analysis

All statistical tests used are detailed in the figure legends. Data in all figures are shown as mean ± SEM. Statistical significance was evaluated using the method indicated in the figure legends.

## Data Availability

Data reported in this paper will be shared by the corresponding author upon request.

## Acknowledgments

We thank Drs. Nieng Yan at Tsinghua University and Zhuo Huang at Peking University for sharing the Na_v_ plasmids; Drs. Xueqin Jin and Jiaofeng Chen at Tsinghua University for technical help of patch-clamp recording of Na_v_; Dr. Xinzhong Dong at the Johns Hopkins University for sharing the Pirt-Cre mice; Dr. Juanjuan Du at Tsinghua University for sharing the anti-Na_v_1.7 antibody; Dr. Zai Chang and other staff at the animal facility center of Tsinghua University for maintaining mice. This work was supported by grant numbers 32425003, 2021ZD0203301, 32130049, 32021002, 31825014 from either the National Natural Science Foundation of China or the National Key R&D Program of China, the New Cornerstone Investigator Program, the Beijing Outstanding Young Scientist Program grant, the Fundamental and Interdisciplinary Disciplines Breakthrough Plan of the Ministry of Education of China (JYB2025XDXM601), the Open Research Fund of Beijing Advanced Center of RNA Biology (BEACON) at Peking University to B.X.

## Author Contributions

B.X. conceived, directed and obtained funding for the study; Y.J. and W.H. contributed equally to this work; Y.J. carried out the bulk experiments related to characterizations of Creld1; W.H. established the postnatal knockout method and did the behavioral genomic screening of the targeted genes; H.Y. did biochemical characterizations of the Creld1-Na_v_ interaction; Y.C. did electrophysiological characterizations of the Creld1-Na_v_ interaction; C.B. did electrophysiological recordings of mechanically activated currents of DRG neurons and helped virus injection; T.Y. helped behavioral experiments; S.W. helped data analysis of DRG single-cell RNAseq data; H.Y. did HCR staining; B.X., Y.J. and W.H. wrote the manuscript; All authors read and edit the manuscript.

## Author Information

The authors declare no competing financial interests. Correspondence and requests for materials should be addressed to B.X.

## Extended Data Figures and Legends

**Extended Data Fig. 1.**
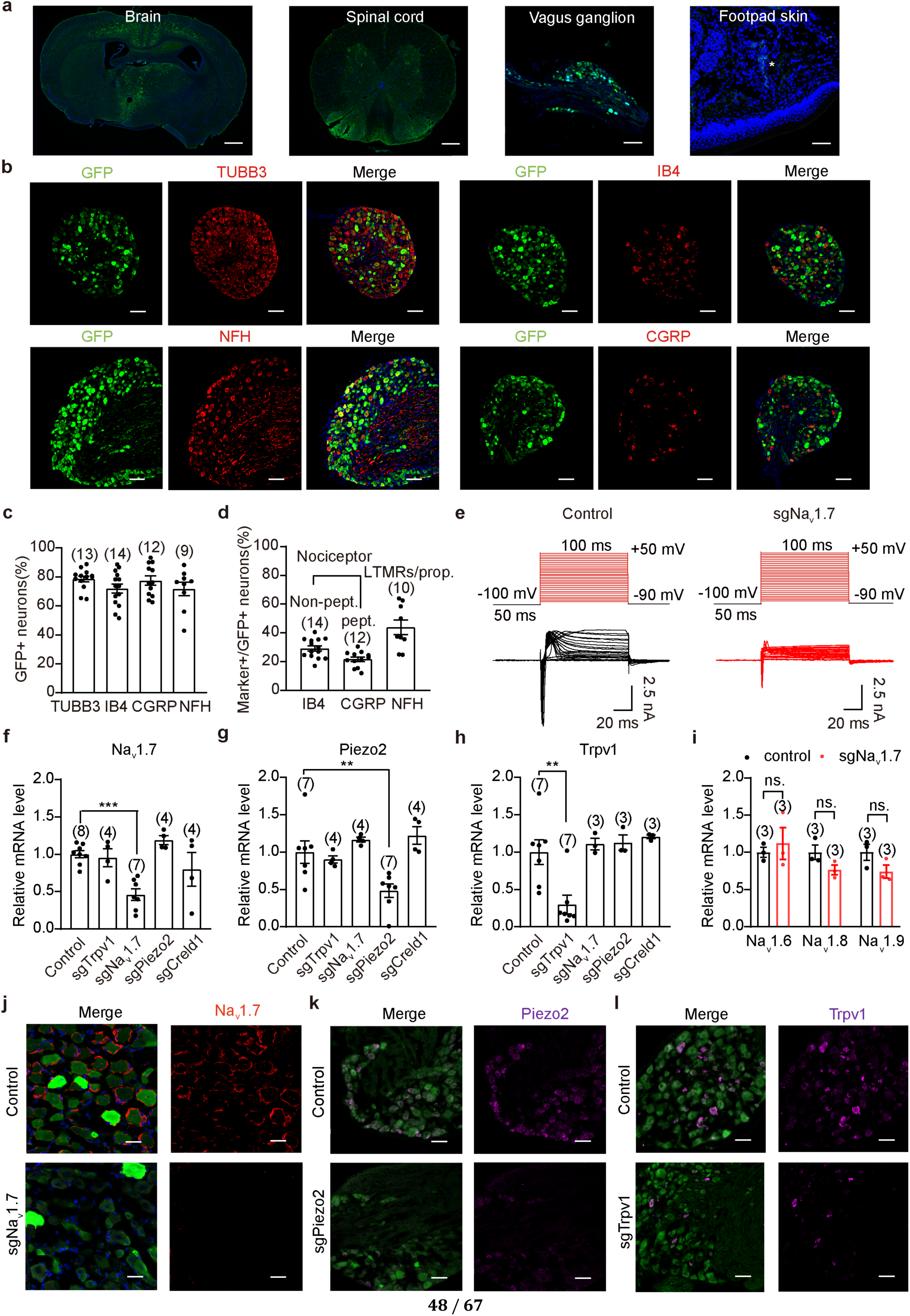
Characterization of AAV9-mediated sgRNA delivery and postnatal knockout of DRG-expressed genes. **a**, Representative immunofluorescence images of brain, spinal cord, vagus ganglion and footpad skin from control mice stained with antibodies against GFP (green) and DAPI. Scale bars, 500 μm for brain; 200 μm for spinal cord; 50 μm for vagus ganglion and footpad skin. **b**, Representative immunofluorescence images of DRG sections from control mice stained with antibodies against GFP (green) and markers for neuronal subpopulations: TUBB3 (pan-neuronal, red), IB4 (non-peptidergic nociceptors, red), NFH (myelinated neurons, red), and CGRP (peptidergic nociceptors, red). Scale bars, 50 μm. **c**, Percentage of GFP-positive neurons in the distinct subsets of DRG neurons classified by the indicated molecular markers. DRG sections were co-stained with anti-GFP antibody and the indicated markers specific for neuronal subpopulations. Each dot represents a DRG section. **d**, Data from panel **c** presented as the percentage of GFP^+^ neurons that co-express a given marker. **e**, Representative TTX-sensitive (TTX-S) sodium currents of control and sgNa_v_1.7 DRG neurons. **f-h**, Scatterplot of the relative mRNA level of *Na_v_1.7*(**f**), *Piezo2* (**g**)and *Trpv1* (**h**) in the indicated sgRNA infected DRG neurons. **i**, Scatterplot of the relative mRNA level of *Na_v_1.6*, *Na_v_1.8* and *Na_v_1.8* in control mice and sgNa_v_1.7 mice. **j**, Immunofluorescent staining of either Na_v_1.7 from the indicated DRG sections. Scale bar, 20 μm. **k** and **l**, HCR (hybridization chain reaction) staining of the Piezo2 mRNA(**k**) and the Trpv1 mRNA (**l**) from the indicated DRG sections. Scale bar, 40 μm. Data are presented as means ± SEM; sample sizes are indicated. One-way ANOVA with Bonferroni’s multiple comparisons test for **f**-**h**; statistical significance is determined by unpaired Student’s *t*-test for **i** (control vs. sgNa_v_1.7 mice), \*\**P* < 0.01, \*\*\**P* < 0.001.

**Extended Data Fig. 2.**
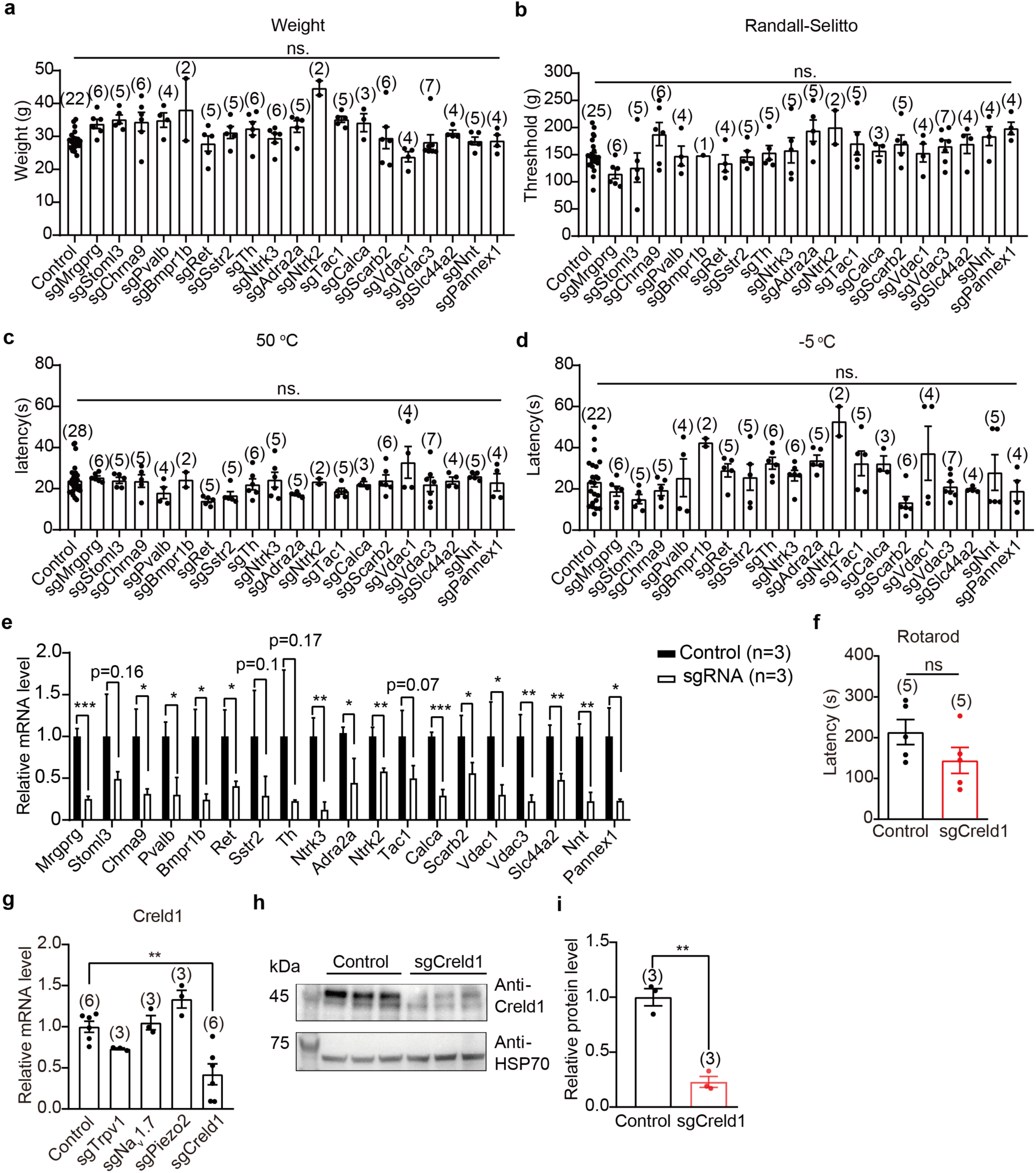
Behavioral assessment of somatosensory functions of mice with postnatal knockout of the targeted genes expressed in DRG neurons. **a**, Scatterplot of body weight. **b**, Scatterplot of the threshold in response to the Randall-Selitto test. **c**, Scatterplot of the paw withdrawal latency in response to the hot plate test. **d**, Scatterplot of the paw withdrawal latency in response to the cold plate test. **e**, Scatterplot of the relative mRNA level of candidate genes. **f**, Scatterplot of the latency to fall in the Rotarod test. **g**, Scatterplot of the relative mRNA level of Creld1 in the indicated sgRNA infected DRG neurons. **h**, Representative images showing western blot of DRG lysates from control and sgCreld1 mice with the antibodies against Creld1 and HSP70. **i**, Scatterplot of normalized protein levels of Creld1. Data are presented as means ± SEM; sample sizes are indicated. Statistical significance is determined by one-way ANOVA with Bonferroni’s multiple comparisons test for **a**-**d, g**. Unpaired Student’s *t*-test for **e**, **f**, **i**, \**P* < 0.05, \*\**P* < 0.01.

**Extended Data Fig. 3.**
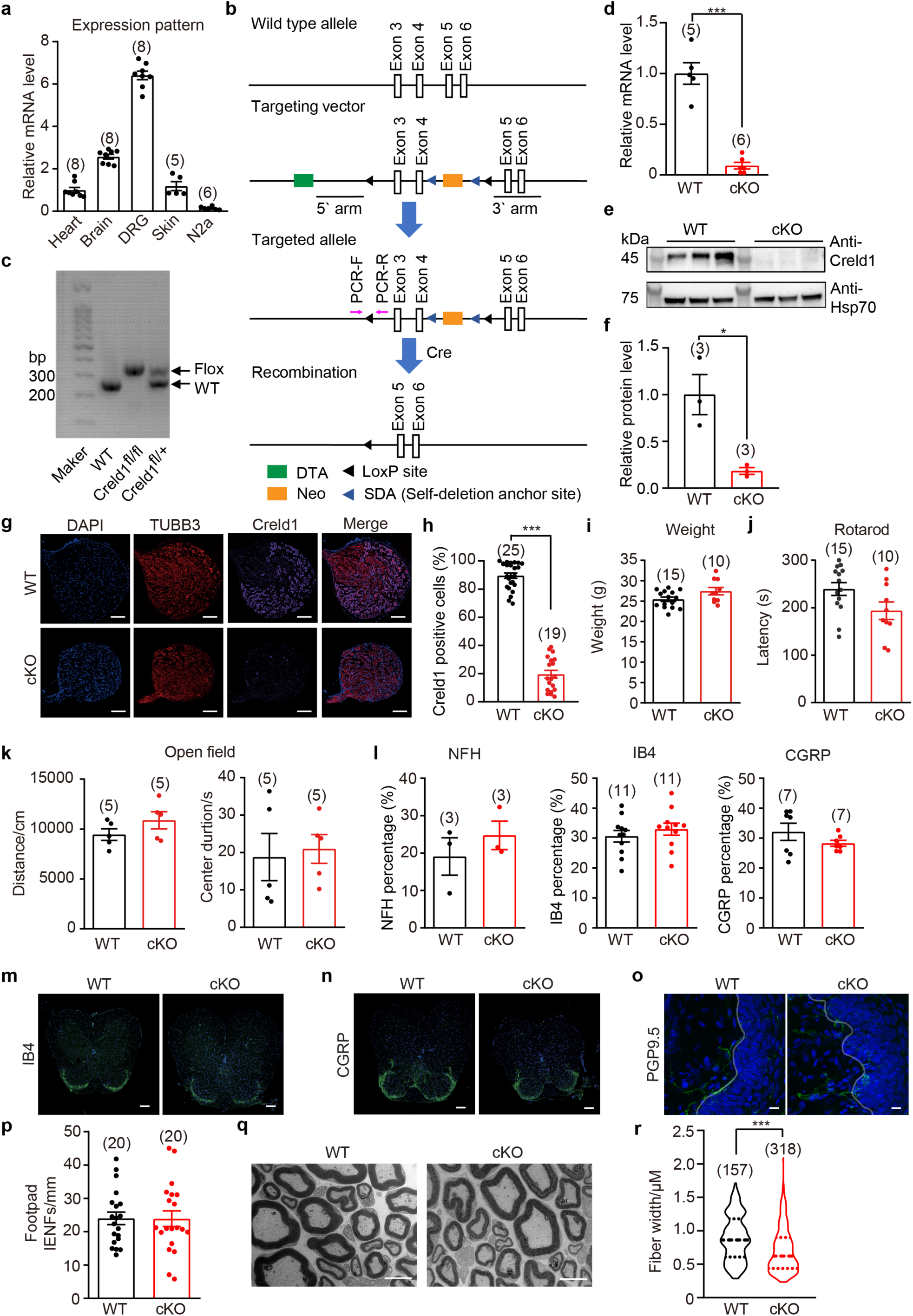
Generation and characterization of the Creld1 conditional knockout mice. **a**, Expression pattern of Creld1 in various tissues. **b**, Construct design for generating the Creld1 floxed mice. Two loxP sites flanking exons 3-4 of the Creld1 locus were inserted via homologous recombination in embryonic stem (ES) cells using a targeting vector with 5’ and 3’ homologous arms. The Creld1^fl/fl^ mice were crossed with tissue-specific Cre line (e.g., Advillin-CreERT2, enabling tamoxifen-inducible, DRG-specific recombination) mice to generate the Creld1^fl/fl^ (WT) and Advillin-CreERT2^/^Creld1^fl/fl^ (tamoxifen treated, cKO) littermates. **c**, Genotyping of the indicated mice using PCR. **d**, Normalized Creld1 mRNA level in DRGs from WT and cKO mice. **e**, Western blots showing Creld1 protein levels in DRG lysates from WT and cKO mice. **f**, Normalized Creld1 protein level of DRGs from WT and cKO mice. **g**, Representative images of immunostaining of DRG sections from WT and cKO mice. Scale bar: 50 μm. **h**, Scatterplot of the percentage of Creld1 positive cells in WT and cKO DRG neurons. **i**, Body weight analysis of WT and cKO mice. **j**, Scatterplot of the latency to fall during the Rotarod test. **k**, Scatterplots showing distance traveled (left panel) and center time duration (right panel) in the open field for WT and cKO mice. **i**, Proportion of distinct DRG neuronal subsets classified by NFH, IB4 and CGRP in WT and cKO mice. **m**, Representative images showing co-immunostaining for DAPI and IB4 of spinal cord sections from WT and cKO mice. Scale bars, 100 μm. **n**, Representative images showing co-immunostaining for DAPI and CGRP of spinal cord sections from WT and cKO mice. Scale bars, 100 μm. **o**, Representative images of immunostaining for PGP9.5 and DAPI of footpad sections from WT and cKO mice. Scale bar: 10 μm. **p**, Quantification of PGP9.5-positive intra-epidermal nerve fibers (IENFs) of footpad from WT and cKO mice. The grey line represents the division between the epidermis and dermis. **q**, Ultrastructural analysis of sciatic nerves from WT and cKO mice. Scale bar, 5 μm. **r**, Quantification for fiber width of sciatic nerves from WT and cKO mice. Data are presented as means ± SEM; sample sizes are indicated. Statistical significance is determined by unpaired Student’s *t*-test for **d**, **f**, **h-l**, **p**, **r**, \**P* < 0.05, \*\*\**P* < 0.001.

**Extended Data Fig. 4.**
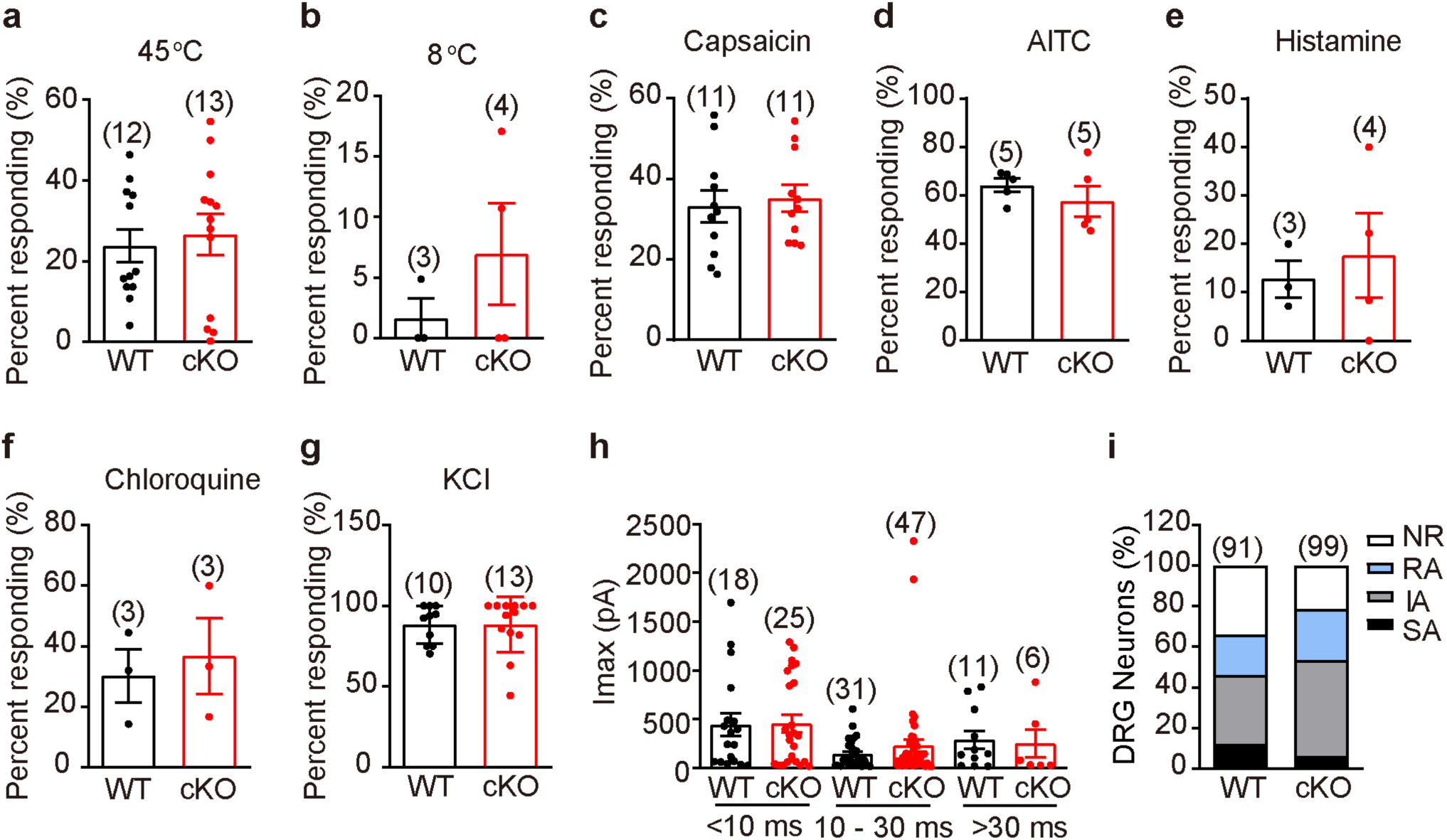
Sensory transduction of WT and cKO DRG neurons. **a**-**g**, Scatterplot of the percentage of WT and cKO DRG neurons responding to heat (**a**), cold (**b**), 10 μM capsaicin (**c**), 100 μM AITC (**d**), 50 μM histamine (**e**), 1 mM chloroquine (**f**) and 50 mM KCl (**g**). **h**, Scatterplot of the maximum current amplitude of the indicated type of currents recorded from WT and cKO DRG neurons. **i**, Proportion of WT and cKO DRG neurons showing the indicated responding properties. Data are presented as means ± SEM; sample sizes are indicated. Statistical significance is determined by unpaired Student’s *t*-test for **a-g**.

**Extended Data Fig. 5.**
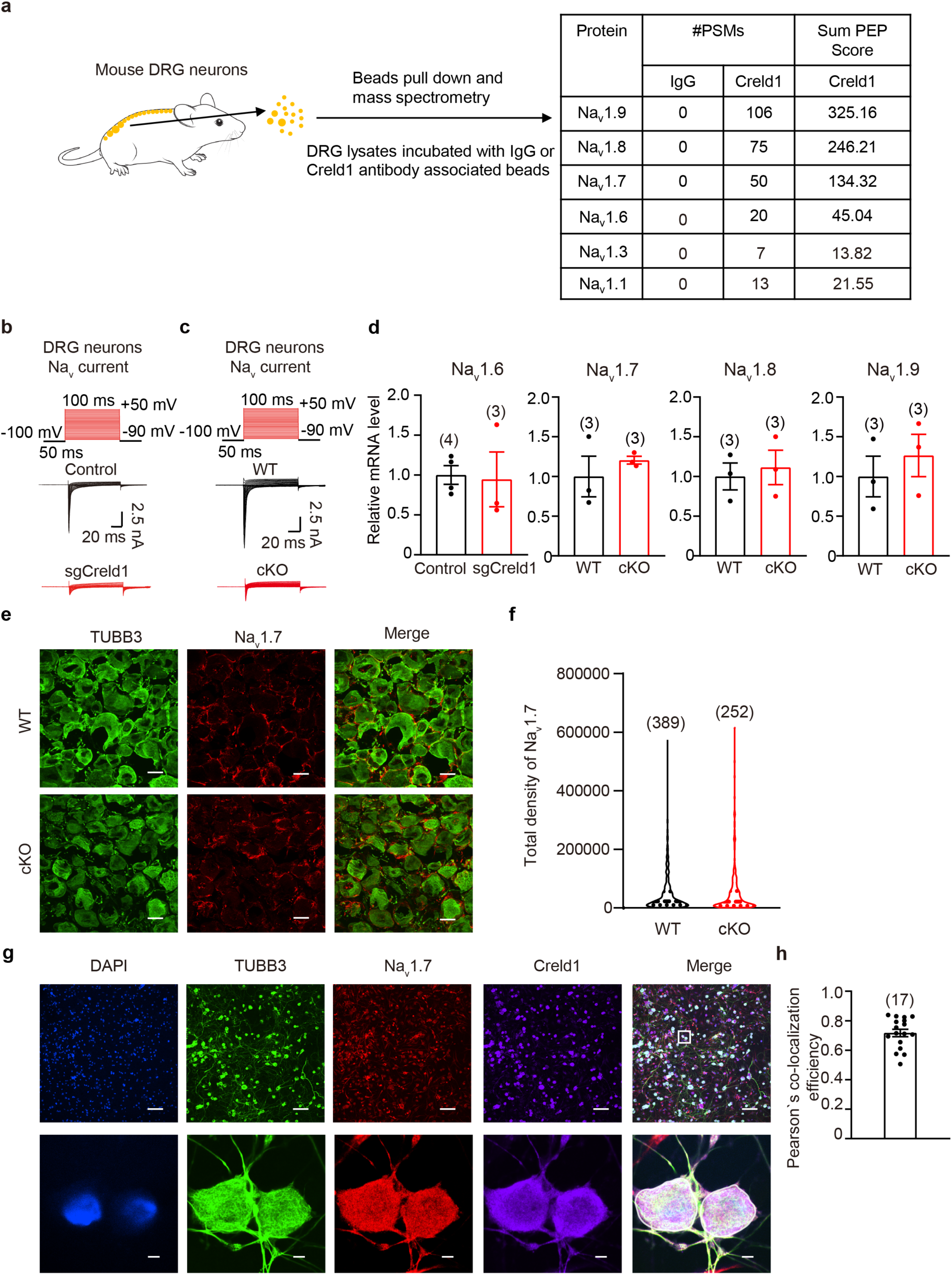
Identification of voltage-gated sodium channels as Creld1-interacting proteins. **a**, Schematic illustration for identification of Creld1-interacting protein in DRGs by immunoprecipitation and mass spectrometry. **b**, Representative Na_v_ currents in DRG neurons from control and sgCreld1 mice. **c**, Representative Na_v_ currents recorded in DRG neurons derived from WT and cKO mice. **d**, Scatterplot of normalized mRNA levels of the indicated Na_v_ isoforms expressed in WT and cKO DRGs. **e**, Immunofluorescent staining of either Na_v_1.7 and TUBB3 from the indicated DRG sections. Scale bar, 20 μm. **f**, Total density of Na_v_1.7 expression of DRG neurons from WT (389 cells) and cKO (252 cells). **g**, Immunofluorescent staining of endogenous Creld1 and Na_v_1.7 in DRG neurons after 36 h in culture using the anti-Creld1, anti-Na_v_1.7 and anti-TUBB3 antibody. Scale bars, 100 μm (top); 5 μm (bottom). **h**, Pearson’s co-localization efficiency analysis for Na_v_1.7 and Creld1 located in DRG neurons after 36 h in culture. The white circle illustrates the region of interest (ROI) used for analyzing the immunofluorescence staining of Na_v_1.7 and Creld1. Each dot represents an individual cell (17 cells in total).

**Extended Data Fig. 6.**
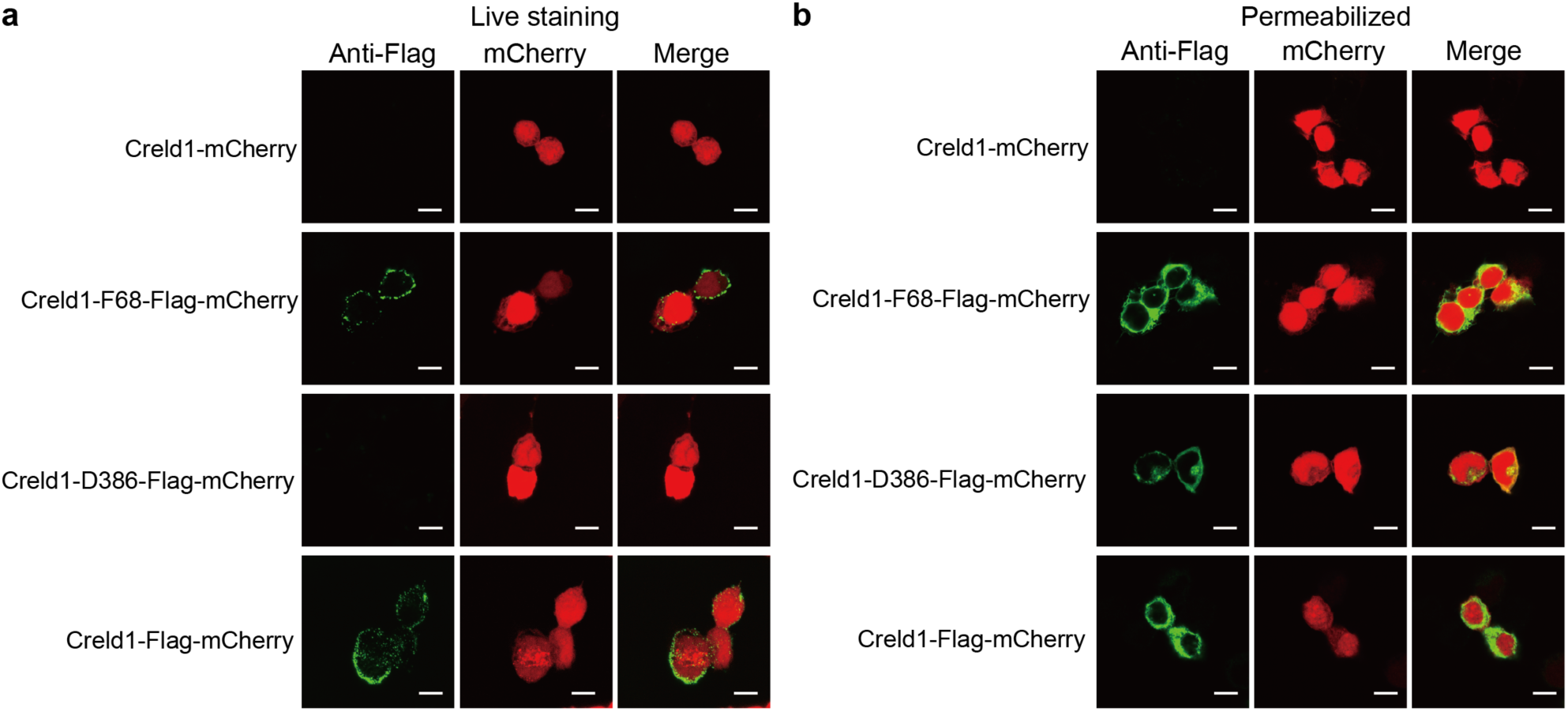
Characterizations of the membrane location and topology of Creld1. Live (**a**) or permeabilized (**b**) immunostaining of Creld1 or Credl1 with the Flag-tag inserted after residue F68, D386 or at the C-terminus of Creld1-IRES-mCherry in HEK293T cells. The mCherry images were taken as control for the transfection of the constructs. Scale bar, 10 μm.

**Extended Data Fig. 7.**
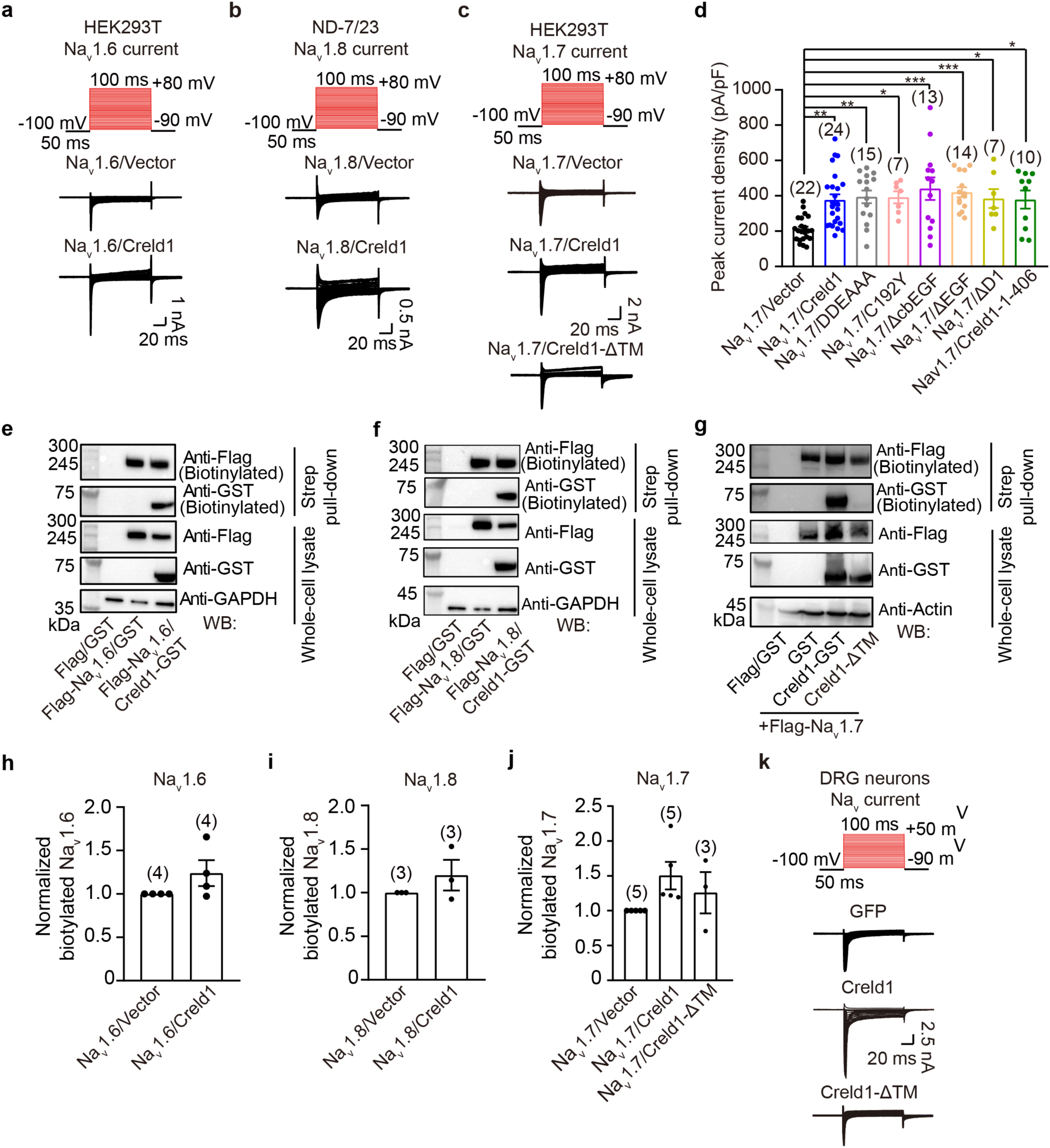
Characterization of the effect of Creld1 on the function and plasma membrane localization of voltage-gated sodium channels. **a-c**, Representative voltage gated sodium currents from the indicated cells transfected with the indicated constructs. **d**, Scatterplot of peak current density of HEK293T cells transfected with the indicated constructs. **e-g**, Western blots of the biotinylated or whole-cell lysate samples derived from HEK293T cells transfected with the indicated constructs. **h-j**, Scatterplot of the normalized biotinylated voltage-gated sodium channel levels in HEK293T cells transfected with the indicated constructs. **k**, Representative voltage gated sodium currents from the indicated DRG neurons. Data are presented as means ± SEM; sample sizes are indicated. One-way ANOVA with Bonferroni’s multiple comparisons test for **d**, or unpaired Student’s *t*-test for **h-j**, \**P* < 0.05, \*\**P* < 0.01, \*\*\**P* < 0.001.

**Extended Data Fig. 8.**
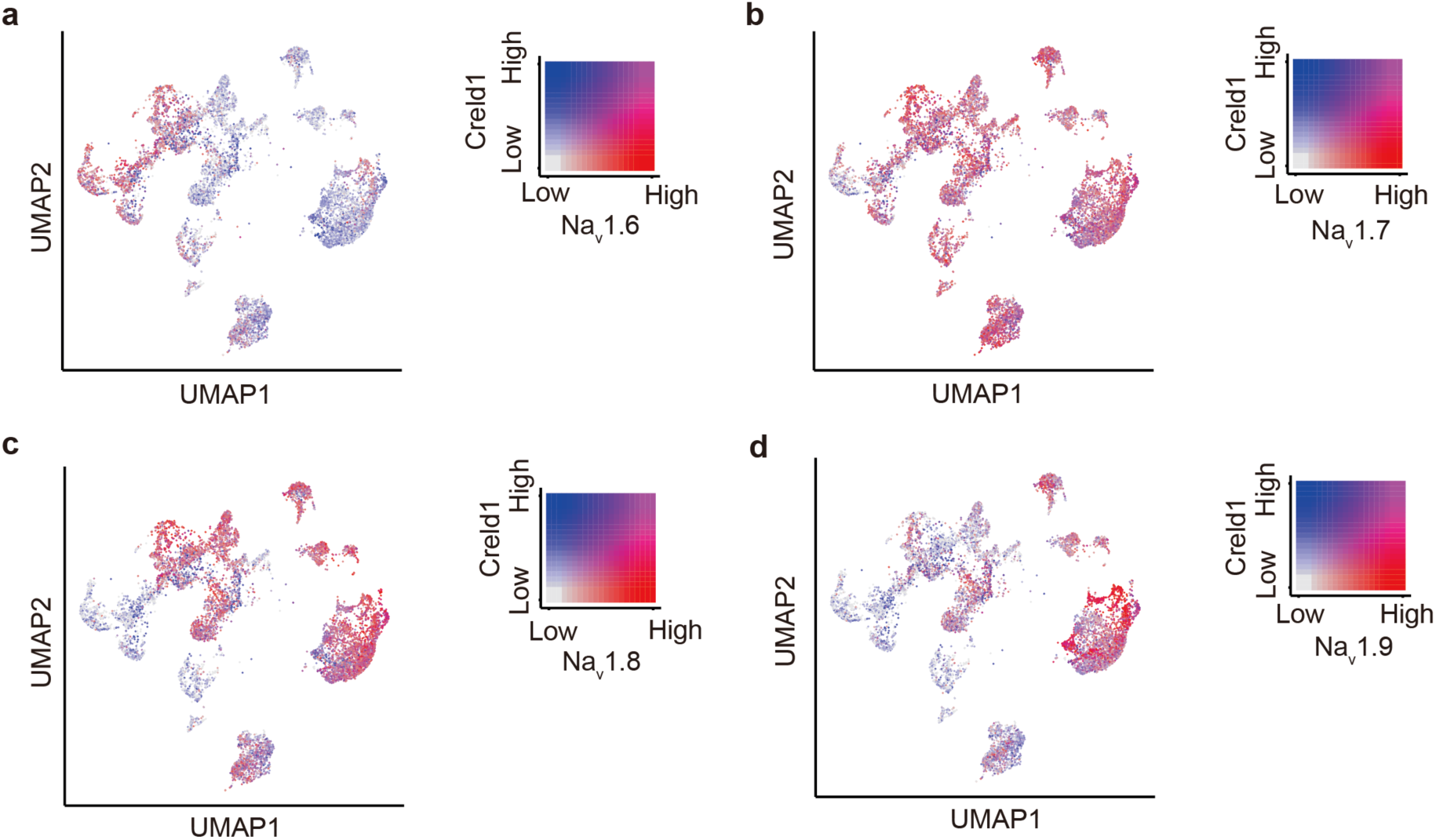
Harmonized human somatosensory neuronal cell atlases for Creld1 and Na_v_. **a**-**d**, Left: UMAP projection of harmonized neuronal atlas, showing expression of Creld1 and Na_v_1.6 (**a**), Na_v_1.7 (**b**), Na_v_1.8 (**c**) or Na_v_1.9 (**d**). Cells are colored according to gene expression: blue for Creld1 and red for Na_v_. Right: Dot plots depicting co-expression patterns of Creld1 and each Na_v_ channel across human DRG neuron subtypes. Color intensity indicates mean log-normalized expression values.

**Extended Data Table 1.**
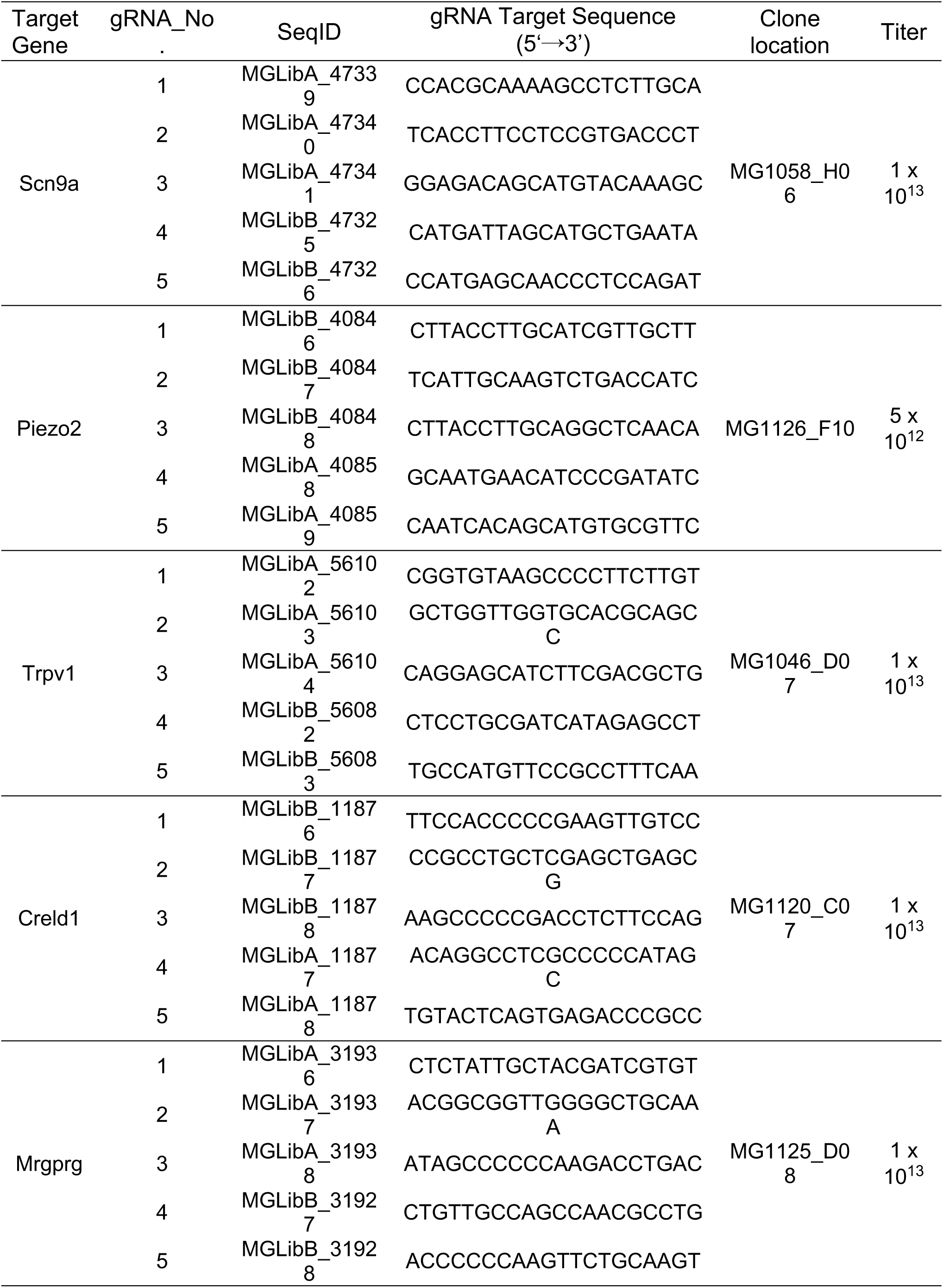
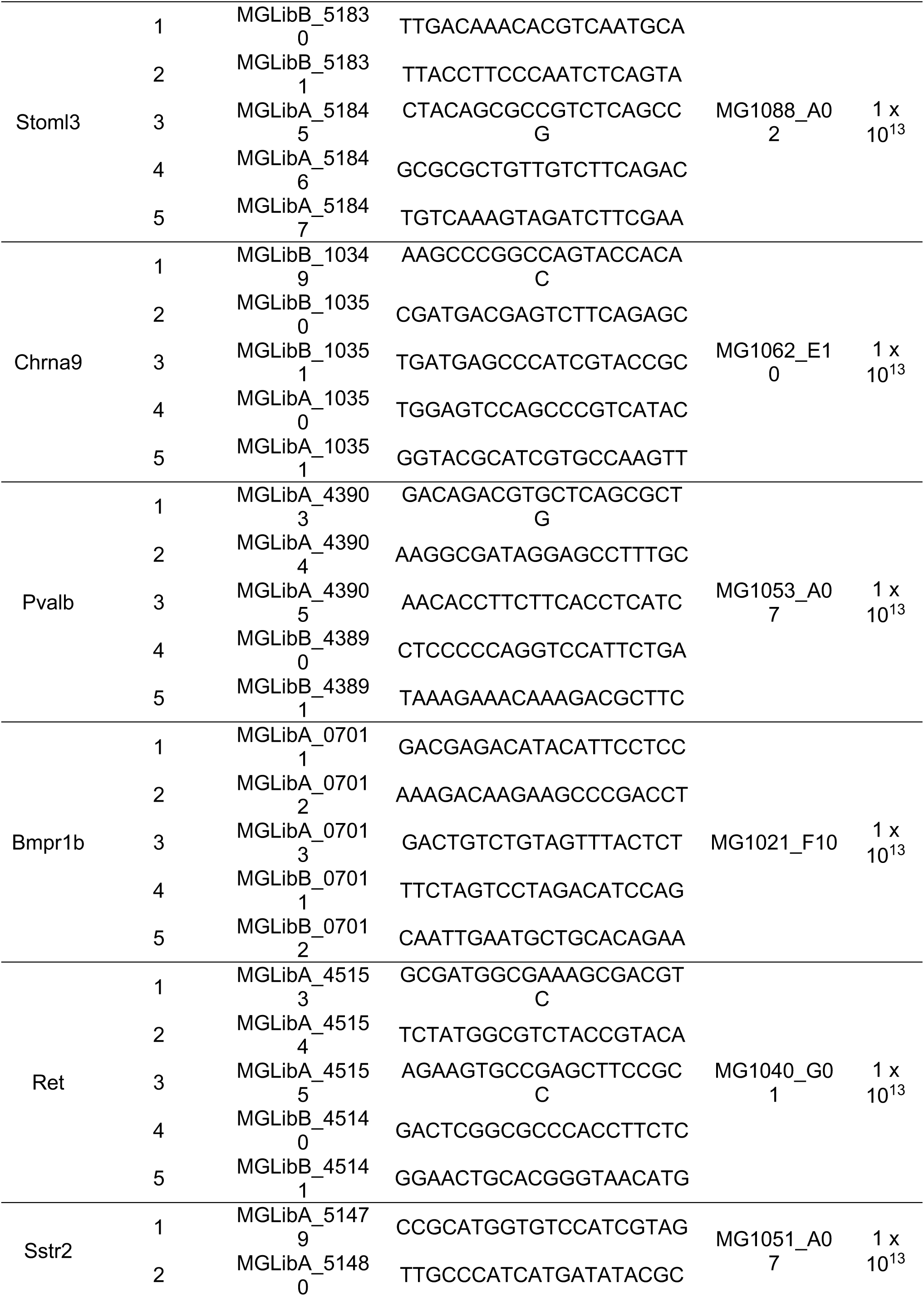
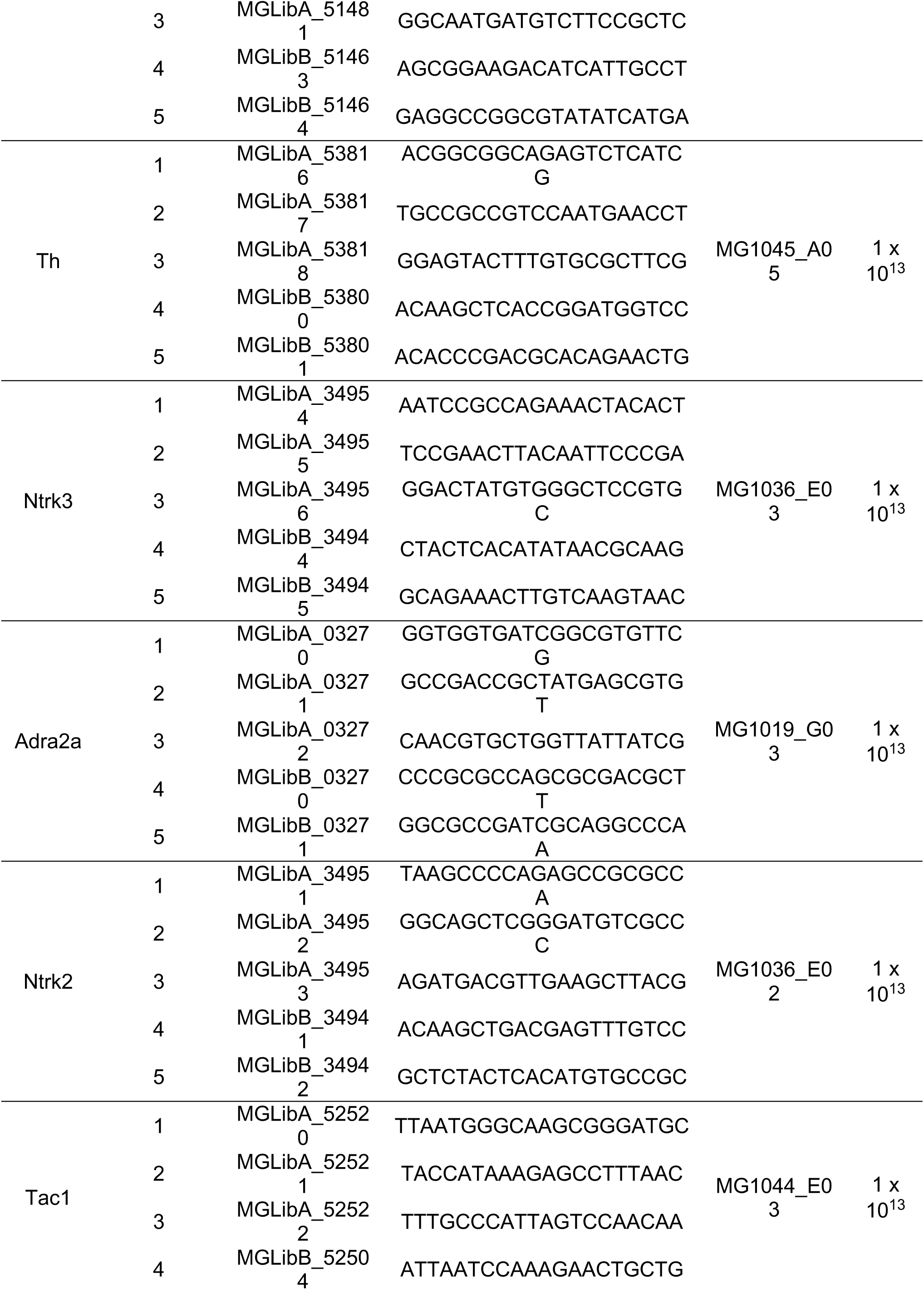
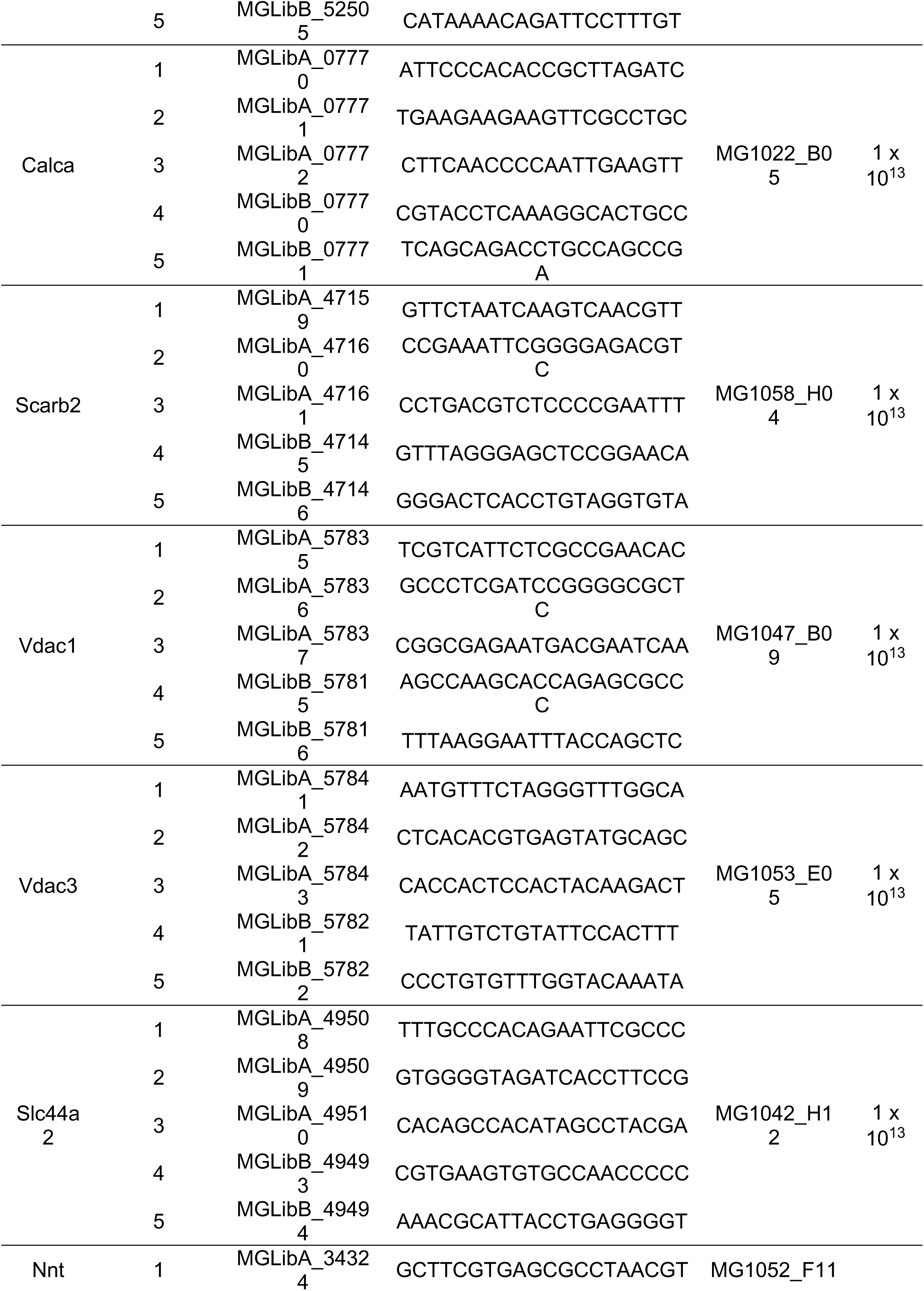
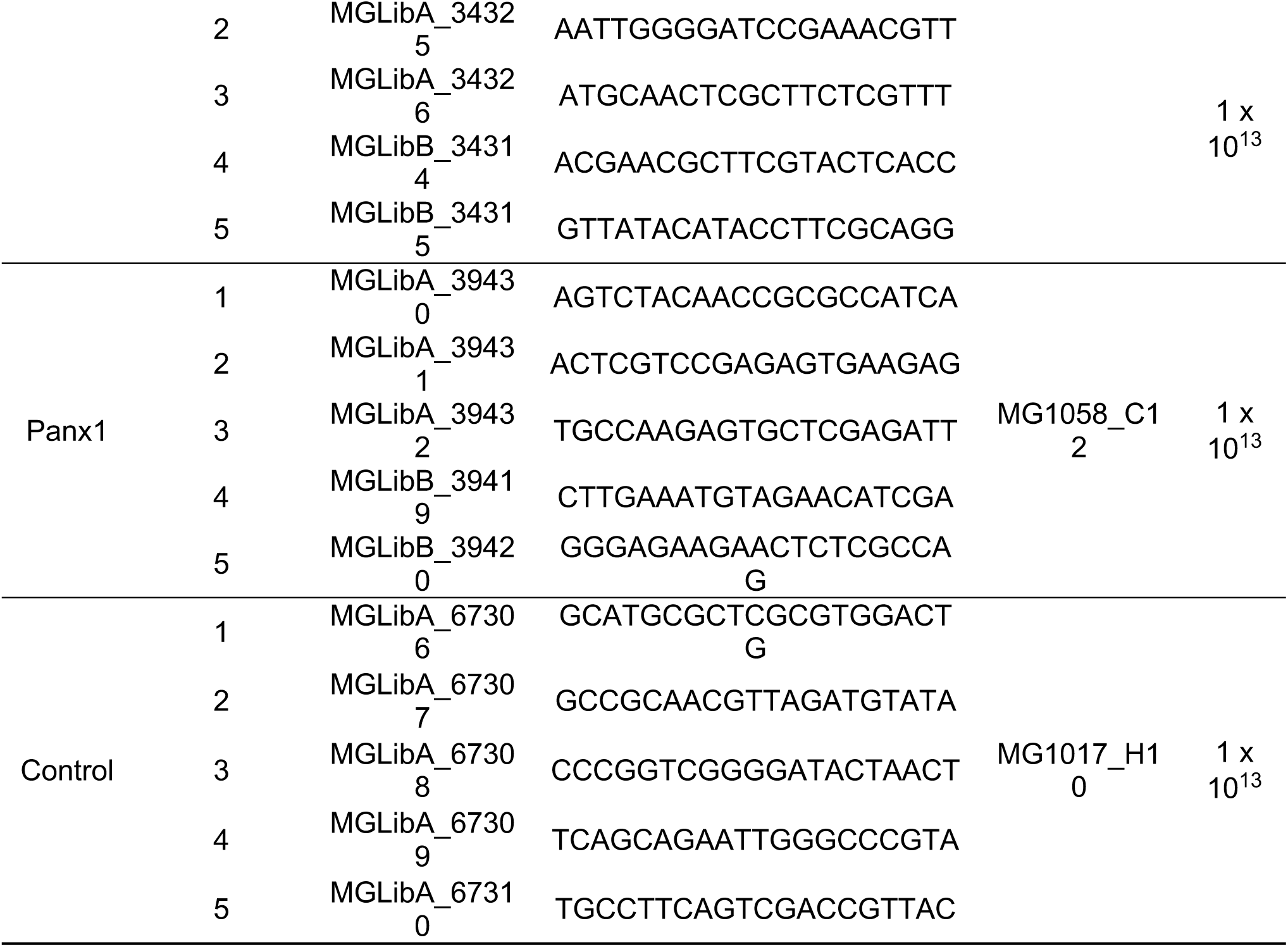

**Extended Data Table 2.**
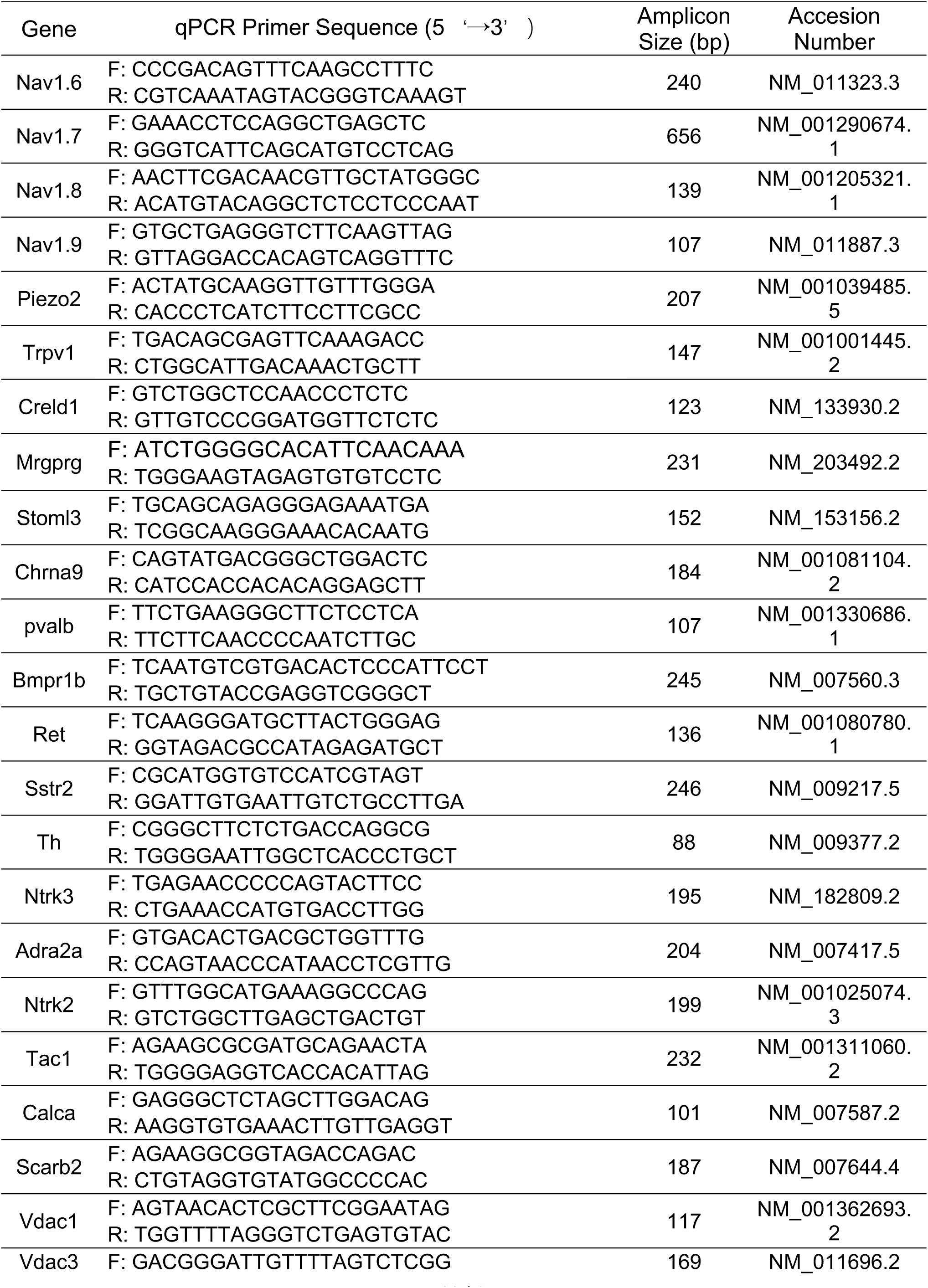
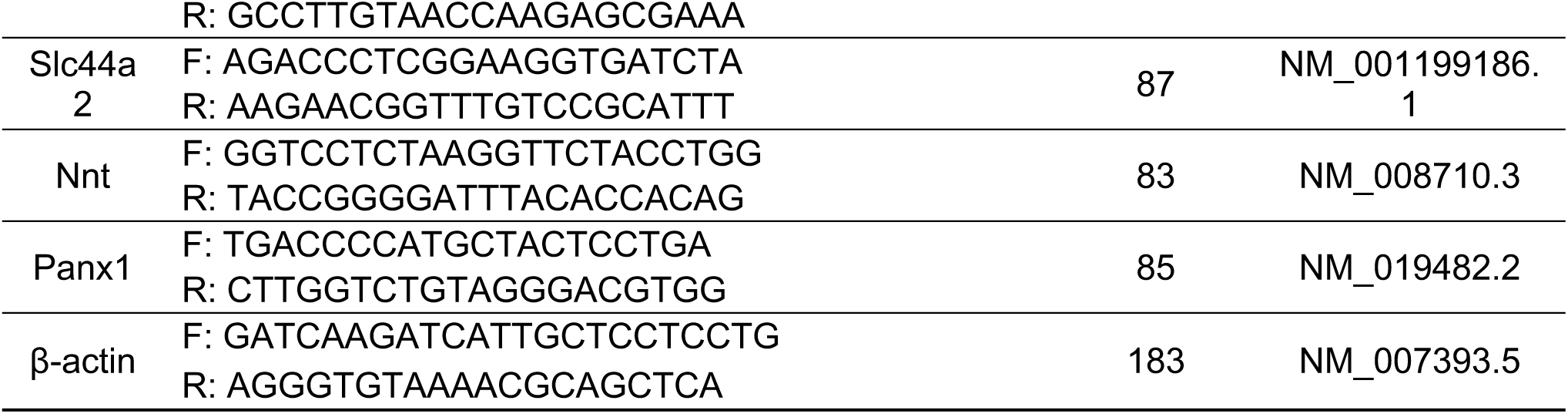

